# CRTC2 regulates plasma cell metabolism and survival

**DOI:** 10.1101/2021.04.14.439620

**Authors:** Jason S Hong, Fasih M Ahsan, Encarnacion Montecino-Rodriguez, Peter D Pioli, Min-sub Lee, Thang L Nguyen, David G Brooks, Justin Golovato, Kayvan R Niazi, Kenneth Dorshkind, Michael A Teitell

## Abstract

Antibody secreting cell (ASC) function and longevity determines the strength and durability of a humoral immune response. Previously, we identified the inactivation of the CREB-regulated transcriptional coactivator-2 (CRTC2) in an *in vitro* B cell differentiation assay that produced functional ASCs. However, the requirement for CRTC2 inactivation in ASC physiology *in vivo* remains unknown. Using transgenic (TG) mice that express a constitutively active form of CRTC2 (*Crtc2-AA*) as an experimental tool, we demonstrate that *Crtc2* repression in plasma cells (PCs) is an intrinsic requirement for ASC metabolic fitness. Sustained CRTC2 activity shortens the survival of splenic and bone marrow PCs, resulting in reduced numbers of long-lived PCs and antibody deficits against T cell dependent and independent antigens, and an acute viral infection. TG PCs resemble short-lived PCs with reductions in glycolysis, oxidative metabolism, spare respiratory capacity, and antibody secretion. Mechanistically, *Crtc2* repression is necessary for the fidelity of PC gene expression and mRNA alternative-splicing programs. Combined, *Crtc2* repression in PCs must occur to support PC metabolism and extend ASC survival during a humoral immune response.

## Introduction

In secondary lymphoid tissues, B cell activation in response to certain antigens can result in the rapid differentiation of transient plasmablasts (PBs) that localize to extrafollicular regions and mature into predominantly short-lived plasma cells (SLPCs). These antibody secreting cells (ASCs) typically produce an initial wave of low-affinity antigen-specific antibodies (MacLennan, Toellner et al., 2003). By comparison, primary follicular B cells activated by a T cell dependent (TD) antigen at the T:B border zone can rapidly expand within B cell follicles to form germinal centers (GCs). GC B cells undergo both extensive proliferation and antigen-driven affinity maturation resulting in the formation of plasma cells (PCs) that secrete high-affinity isotype-switched antibodies. These PCs can remain locally or more typically migrate to the bone marrow and become long-lived PCs (LLPCs) (De Silva & Klein, 2015). Antibody quality and the duration of antibody production are major determinants of protection against target pathogens and maintenance of long-term humoral immunity (Amanna, Carlson et al., 2007). Unfortunately, the production of pathogenic PCs can also generate autoantibodies that participate in a range of autoimmune disorders (Brink & Phan, 2018). Identifying factors that regulate PC differentiation, function, and survival could improve vaccine design and uncover potential strategies to eliminate the production and/or maintenance of pathogenic PCs.

To generate ASCs, extensive epigenetic, transcriptional, and metabolic remodeling must occur to terminate B cell and GC B cell specification and maintenance programs, coupled with induction of ASC differentiation (Barwick, Scharer et al., 2016, Boothby & Rickert, 2017, Nutt, Hodgkin et al., 2015). Differing in their ontogeny, localization, function, and lifespan, ASCs are present as a heterogenous and diverse population. RNA expression profiling of splenic PCs that are enriched for SLPCs and bone marrow PCs that are enriched for LLPCs failed to identify wholesale transcriptional changes that could explain phenotypic differences in SLPC and LLPC longevity (Lam, Jash et al., 2018). Instead, LLPCs were shown to develop a unique metabolic program that includes enhanced glucose import and increased pyruvate utilization for mitochondrial oxidative metabolism during times of stress (Lam, Becker et al., 2016). How this metabolic program is initiated and maintained in LLPCs is largely unknown. The lack of candidate gene expression differences between SLPCs and LLPCs that underlie differences in survival or longevity may suggest non-transcriptional or extrinsic factors. However, because of heterogeneity in the ASC population, meaningful mRNA transcript differences may actually exist that are muted within this heterogeneity and therefore overlooked.

Identified over a decade ago, a family of three cAMP response element binding protein (CREB)-regulated transcription coactivators (CRTC1 – 3) control gene expression and several additional biological processes (Altarejos & Montminy, 2011, Conkright, Canettieri et al., 2003, Tasoulas, Rodon et al., 2019). Extracellular and intracellular signals phosphorylate CRTC family proteins, which influences their subcellular localization and activity (Jansson, Ng et al., 2008, Koo, Flechner et al., 2005). CRTC proteins lacking phosphorylation at specific residues localize in the nucleus where they can bind to CREB and additional bZIP transcription factors to regulate gene expression for processes such as energy homeostasis, metabolism, and lifespan (Altarejos & Montminy, 2011, Tasoulas et al., 2019). Previously, we showed that CRTC2 phosphorylation in an *in vitro* B cell differentiation model caused CRTC2 inactivation by cytoplasmic re-localization, which triggered B cells to lose their GC-like identity and differentiate into ASCs (Sherman, Kuraishy et al., 2010). Using isolated B cells, DNA damage during *immunoglobulin* (*Ig*) class switch recombination (CSR) initiated a signaling cascade involving ATM, LKB1, and an AMPK family member protein that phosphorylated and inactivated CRTC2 to down-regulate its gene network. Failed CRTC2 inactivation caused GC B cell biomarkers to remain elevated and inhibited ASC biomarker acquisition and antibody secretion (Sherman et al., 2010). However, it remains unknown when CRTC2 inactivation is critical for ASC formation because activation, proliferation, and PC differentiation are plausible regulatory nodes. Additionally, whether CRTC2 plays a critical role in ASC formation *in vivo* has not been investigated.

To examine CRTC2 in B cell physiology, we generated transgenic (TG) mice that express constitutively active, nucleus-localized CRTC2 throughout B cell development. Here, we report on this unique CRTC2 TG mouse model and the unexpected result that CRTC2 inactivation in PCs is required for ASC survival, yielding at least a partial mechanism to explain the longevity of the humoral immune response.

## Results

### *Crtc2* expression in B cells is repressed during PC differentiation

Initially, we examined the expression of CRTC2 in naïve splenic B cells from wild-type (WT) C57BL/6 mice stimulated with CD40L and IL-4, conditions that resemble a TD immune response (Hasbold, Corcoran et al., 2004, Maliszewski, Grabstein et al., 1993). At early time points post-stimulation, CRTC2 lacked phosphorylation and localized in the nucleus, where it can influence target gene expression (Altarejos & Montminy, 2011, Tasoulas et al., 2019). By 24h post-stimulation, approximately half of the CRTC2 protein was phosphorylated, indicated by the emergence of an upper band on western blot (Dentin, Liu et al., 2007, Koo et al., 2005, Screaton, Conkright et al., 2004), followed by a progressive reduction of total CRTC2 protein through day 5 of stimulation (Fig 1A). Phosphorylation at S171 and/or S275 by AMPK family member kinases (Jansson et al., 2008, Screaton et al., 2004) is the source for CRTC2 inactivation with cytoplasmic re-localization. Consistent with a previously identified signal transduction pathway that inactivated CRTC2 in human GC B cells (Sherman et al., 2010), ATM activation by S1981 phosphorylation, and LKB1 activation, shown with phospho-ATM/ATR substrate antibody detection of immunoprecipitated LKB1, preceded and coincided with CRTC2 inactivation by phosphorylation at S171 (Fig 1A). Thus, CRTC2 inactivation by S171 phosphorylation occurs by a conserved signal transduction mechanism in activated human and mouse B cells that were stimulated to differentiate. There is also a progressive reduction in total CRTC2 protein over time.

**Figure 1.**
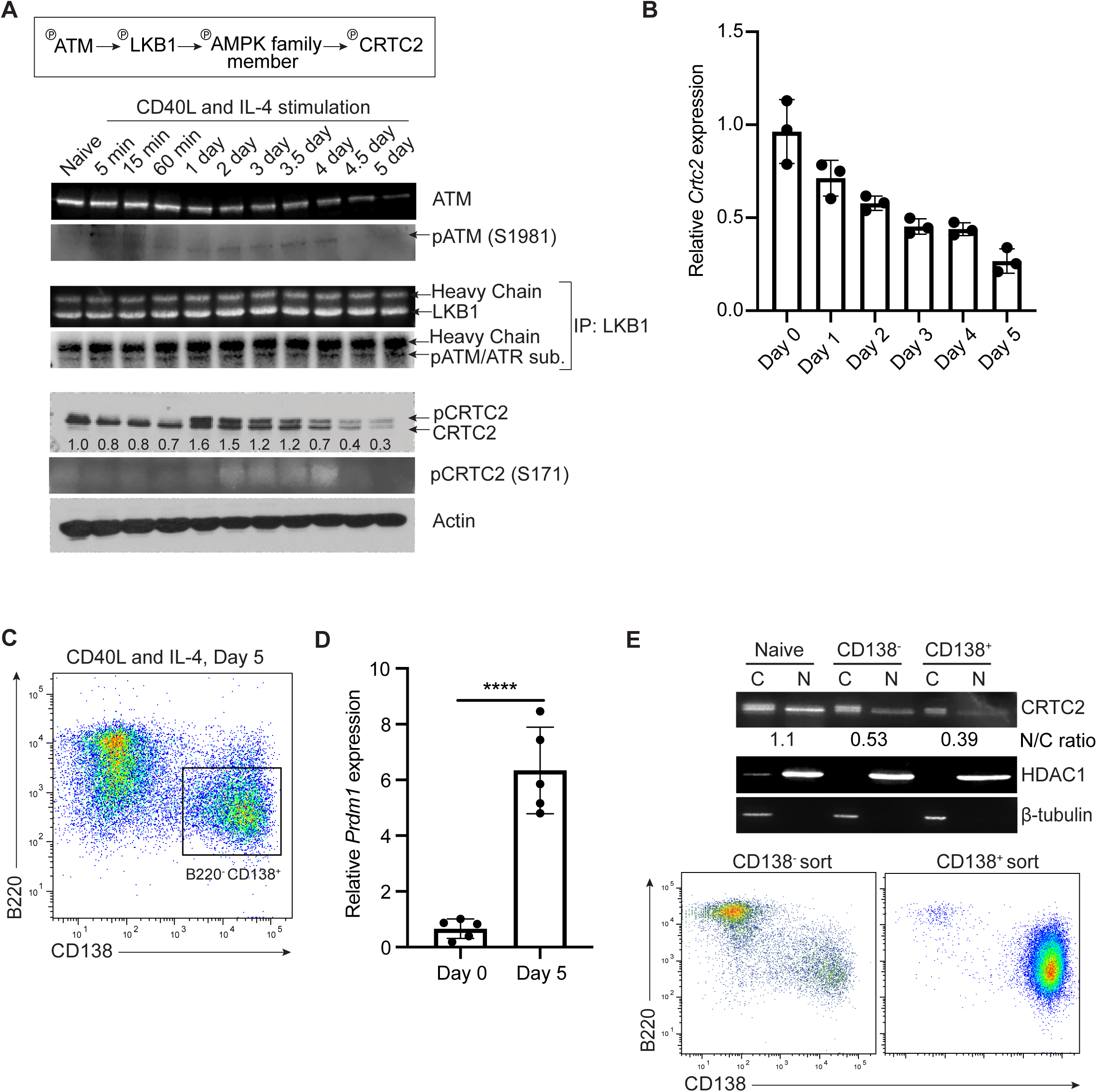
CRTC2 expression in B cells is repressed during PC differentiation. A. Schematic of a signaling pathway that inactivates CRTC2 (top). Immunoblot of a representative time-course analysis of unphosphorylated and phosphorylated ATM, LKB1, CRTC2 protein expression in naïve splenic B cells stimulated with CD40L and IL-4 over 5 days. Actin was used as a loading control. ATM phosphorylation of LKB1 was assessed by LKB1 immunoprecipitation and phospho-(S/T) ATM/ATR substrate immunoblotting. Numbers indicate CRTC2 band intensities normalized to actin and were quantified by ImageJ analysis (*n* = 3 WT and 3 TG). B. Time-course analysis of *Crtc2* mRNA expression from naïve splenic B cells stimulated with CD40L and IL-4 by quantitative RT-PCR (*n* = 3 WT and 3 TG). C. Representative flow cytometry plot at day 5 of CD40L and IL-4 stimulation showing PC differentiation (boxed area). D. Quantification of *Prdm1* expression at day 5 of CD40L and IL-4 induced differentiation by quantitative RT-PCR analysis (*n* = 5 WT and 5 TG). E. Representative immunoblot of CRTC2 expression in the cytoplasm (C) and nucleus (N) of naïve, activated (CD138^-^), and PC/PB (CD138^+^) populations. CRTC2 band densitometry was calculated in arbitrary fluorescent units and presented as the nuclear to cytoplasmic (N/C) ratio. HDAC1 and β-tubulin were used as a nuclear and cytoplasmic control respectively (top). Representative flow cytometry plot showing the enrichment of sorted CD138^+^ PCs/PBs at day 5 of *in vitro* differentiation (bottom) (*n* = 3 WT and 3 TG). Data information: In (B,D), data are presented as mean ± SD. *P* values determined by two-tailed, unpaired Student’s *t*-test, \*\*\*\**P ≤* 0.0001.

Consistent with progressively reduced CRTC2 protein, the level of *Crtc2* mRNA transcripts were progressively reduced over the 5 day differentiation time course (Fig 1B). At day 5, when the level of CRTC2 protein was at its lowest, we identified the differentiation of activated B cells into B220^-^ CD138^+^ PCs by cell surface biomarker expression (Fig 1C). PC differentiation was also confirmed by a 6-fold induction of *Prdm1* transcripts, encoding the ASC master regulator protein BLIMP1 (Shaffer, Lin et al., 2002), in the bulk differentiation culture (Fig 1D). To determine the subcellular localization of reduced CRTC2 protein in PCs, CD138^+^ cells were isolated at day 5 of *in vitro* differentiation, followed by cytoplasmic and nuclear fractionation. CD138^+^ cells included mainly PCs with a minor population of B220^+^CD138^+^ PBs. Western blot showed that CRTC2 was predominantly inactive and repressed in PCs/PBs (CD138^+^) compared to naïve and activated (CD138^-^) B cells (Fig 1E). These results showed that CRTC2 inactivation by protein phosphorylation, re-localization, and decreased transcript and protein levels occurs during the course of B cell differentiation, beginning with early B cell activation and progressing through PC formation.

### Generation of an *in vivo* model to study *Crtc2* repression in PCs

To study potential roles for CRTC2 in mature B cells *in vivo*, we generated a TG mouse model that expressed a nucleus-localized, constitutively active form of CRTC2, generated by targeted S171A and S275A amino acid substitutions in the transgene (*Crtc2-AA*) (Jansson et al., 2008, Screaton et al., 2004, Sherman et al., 2010). The *Crtc2-AA* transgene was under *B29* (*Igβ*) minimal promoter and *IgH* intronic enhancer (*Eμ*) control (Fig EV1A). Previously, we showed that this promoter-enhancer combination enabled transgene expression from pro-B through PC stages of B cell differentiation (Hermanson, Eisenberg et al., 1988, Hoyer, French et al., 2002). Three founder mice were generated with a range of *Crtc2-AA* transgene expression. Two founder lines (F2 and F3) reproducibly showed 3-fold higher levels of *Crtc2-AA* expression in the bone marrow than the third founder line (F1) (Fig EV1B). We analyzed bone marrow B lymphocyte development using flow cytometry of the cell surface biomarkers B220, CD43, and IgM. Founder line F1, with modest total (endogenous and transgene) *Crtc2* overexpression in the bone marrow (1.4-fold compared to WT) (Fig 2A), demonstrated normal production of pro-B (B220^+^ IgM^-^ CD43^+^), pre-B (B220^+^ IgM^-^ CD43^-^), immature (B220^+^ IgM^+^), and mature B220^hi^ IgM^+^ B lymphocytes compared to WT mice (Fig 2B). In contrast, founders F2 and F3 demonstrated a significant reduction in bone marrow B lymphopoiesis (Fig EV1C). While interesting, F2 and F3 founder lines were not examined further because of this impairment in early B cell development. F1 lineage mice showed total *Crtc2* expression that was 3.4-fold higher in the spleen compared to WT mice (Fig 2A), and F1 lineage spleens possessed similar total numbers of follicular (B220^+^ CD19^+^ CD23^+^ CD21^-^) and marginal zone (B220^+^ CD19^+^ CD23^-^ CD21^+^) B cells compared to WT controls (Fig 2C). Finally, low level Igβ (encoded by *B29*) expression was reported by others in early thymocyte development (Wang, Diamond et al., 1998). However, total *Crtc2* expression was not significantly increased in F1 thymocytes (Fig 2A) and the T lineage cell subsets in F1 thymus and spleen were similar to WT controls (Fig 2D and E). Thus, B and T cell development appeared similar between F1 TG and WT mice.

**Figure 2.**
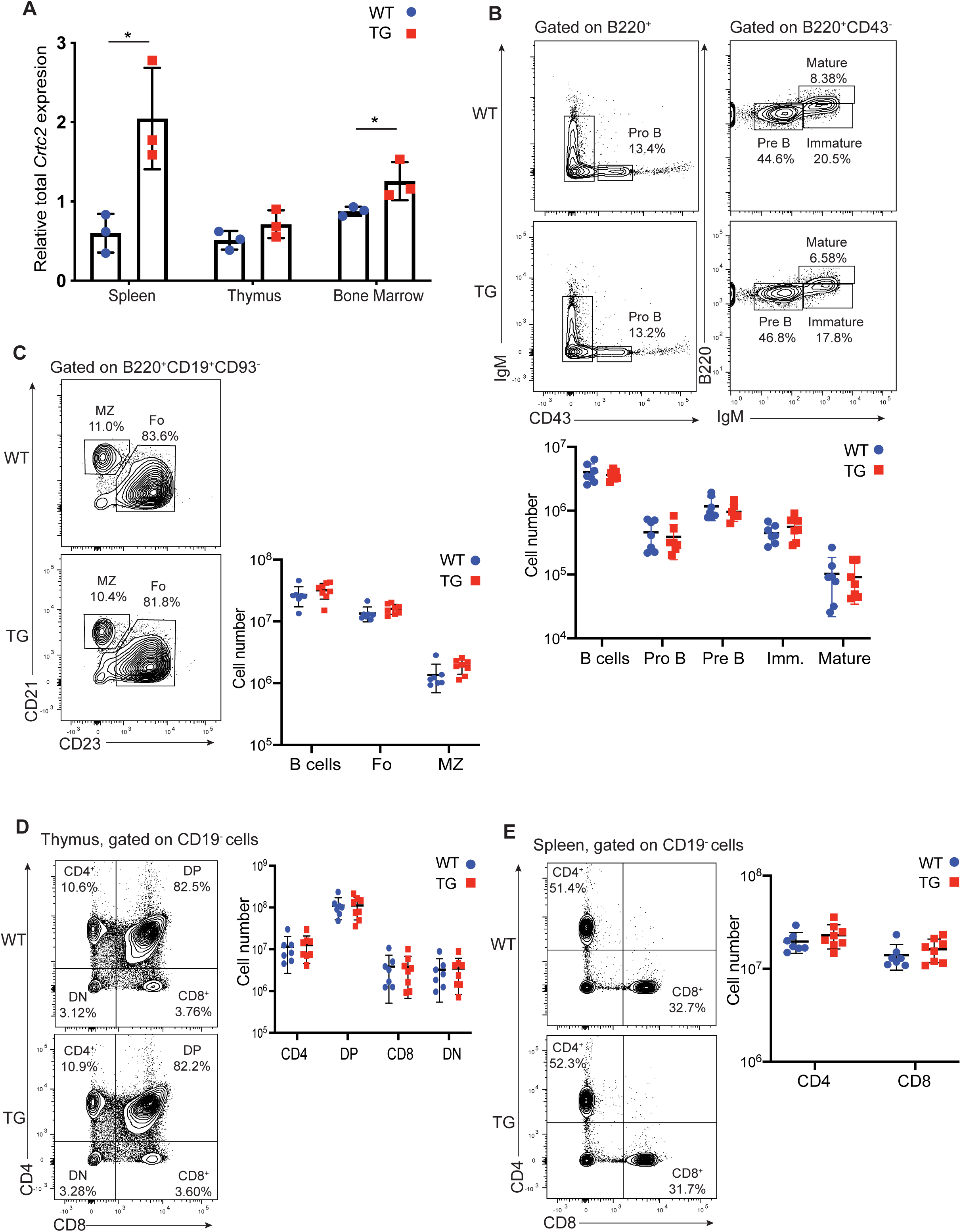
B cell development is unaffected in *Crtc2-AA* F1 TG mice. A. Total *Crtc2* mRNA expression (endogenous and transgene) from mouse cells isolated from the spleen, thymus, and bone marrow by quantitative RT-PCR analysis (*n* = 3 WT and 3 TG). B. Representative flow cytometry plots of B cell populations in the bone marrow of WT and TG mice (top) and quantification of absolute numbers of total, pro-B, pre-B, immature, and mature B cells in the bone marrow (bottom). C. Representative flow cytometry plots of B cell populations in the spleen of WT and TG mice (left) and quantification of absolute numbers of total, FO, and MZ B cells in the spleen (right). D. Representative flow cytometry plots of T cells in the thymus of WT and TG mice (left) and quantification of absolute numbers of subpopulations in the thymus (right). E. Representative flow cytometry plots of T cells in the spleen of WT and TG mice (left) and quantification of absolute numbers of T cell subpopulations in the spleen (right). Each symbol (B-E) represents an individual mouse (*n* = 7 WT and 8 TG). Data information: In (A-E), data are presented as mean ± SD. *P* values determined by two-tailed, unpaired Student’s *t*-test, \**P ≤* 0.05.

To exclude transgene insertion site issues in F1 lineage mice, we performed whole genome sequencing and identified two distinct *Crtc2-AA* genome integration sites. One integration site was in a region of the mouse genome lacking any genes and the second site was in a large intronic region of a gene (*1110051M20Rik*) with no functional annotation (Fig EV1D). This second transgene integration had no effect on the expression of the annotated gene as determined by RNA sequencing (RNA-seq) of TG and WT B cells (Table EV1).

Next, we examined the level of *Crtc2* expression in activated B cells. We stimulated naïve splenic B cells with CD40L, IL-4, and IL-5 for 5 days, with IL-5 added to increase the number of PCs obtained (Hasbold et al., 2004). On day 5, stimulated B cells were sorted into early division (0-3), late division (5 or more), and PC subpopulations based on CFSE dye dilution and expression of the ASC cell surface marker CD138 (Fig EV1E). We further validated the identity of *in vitro* derived and sorted PCs by comparing expression profiles to *in vivo* derived PC gene signatures (Shi, Liao et al., 2015). RNA-seq of naïve B cell controls and sorted, stimulated B cell subpopulations revealed stable endogenous *Crtc2* expression in naïve, early, and late division B cell subsets, along with robust *Crtc2* repression in WT PCs (Fig EV1F). By contrast, *Crtc2-AA* TG B cells showed minimally elevated total *Crtc2* expression in naïve and early division B cell subsets, with 1.4-fold and 4.4-fold increased expression in late division and PC subpopulations, respectively (Fig EV1F). Induced expression of total *Crtc2* in TG PCs is consistent with increased *IgH* intronic enhancer activity in the transgene that is known to elevate *Ig* gene expression at the PC stage of B cell differentiation (Ong, Stevens et al., 1998). These data validate our *Crtc2-AA* TG model that replicates expected transgene expression throughout B cell differentiation and provides sustained, over-expressed *Crtc2-AA* transcripts specifically in PCs, the stage in differentiation where endogenous *Crtc2* is strongly repressed.

### CRTC2 repression enables PC survival and humoral immune responses

Our prior *in vitro* studies showed that aberrantly sustained CRTC2 activity following B cell activation caused a defect in ASC differentiation (Sherman et al., 2010). However, this pure B cell model left open the key question about the main role(s) for CRTC2 repression during an *in vivo* immune response. Analysis of the ASC population in naïve unimmunized mice revealed a significant decrease in the splenic PC population, while the PB population was unaffected in TG mice compared to WT controls (Fig 3A). Further analysis of the serum of naïve mice revealed lower Ig titers in TG compared to WT mice for all isotypes, except for IgA (Fig 3B). Next, we immunized mice with TNP-Ficoll to induce a T cell independent (TI) immune response. TNP-specific IgM titers in the serum were 7-fold lower in TG compared to WT mice 7 days post-immunization (Fig 3C). Mice were also immunized with NP-CGG in alum to induce a TD immune response. At 14 days post-immunization, NP-specific IgM and IgG1 titers were 2.7- and 6-fold lower in the serum of TG compared to WT mice, respectively (Fig 3D). At 35 days, NP-specific serum IgG1 levels remained significantly lower in TG compared to WT mice, revealing that reduced Ig levels are not due to slower ASC differentiation in TG B cells (Fig 3D). The secondary humoral immune response was then assessed at this time point by NP-CGG re-immunization. Indicative of a functional memory B cell pool, both TG and WT mice showed a further increase in IgG1 levels 7 days post re-immunization, although IgG1 remained significantly reduced in TG compared to WT mice (Fig 3D). We also infected mice with the acute Armstrong strain of lymphocytic choriomeningitis virus (LCMV) to analyze the antiviral B cell response. This viral strain produces a vigorous T cell response that clears the infection 8 – 10 days post-infection but also elicits a robust TD B cell response, which generates LLPCs (Ahmed, Salmi et al., 1984, Hao, Li et al., 2018). Analysis of the spleen 8 days post-infection revealed a sharp reduction in the percentage and total number of PCs in TG mice (Fig 3E), along with a marked reduction in Ig titers for both total and LCMV-specific IgG compared to WT control mice (Fig 3F and G).

**Figure 3.**
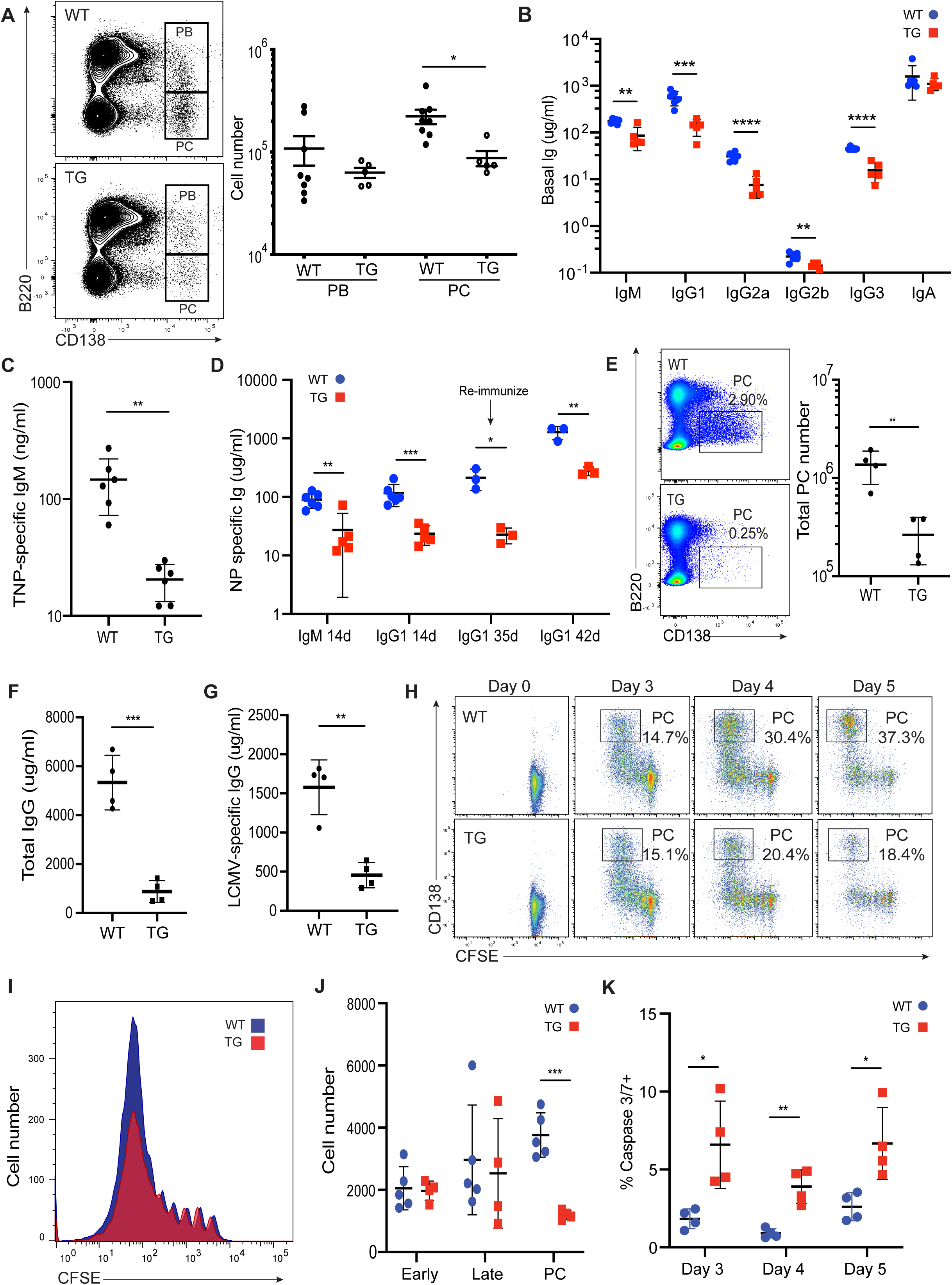
Impaired humoral immune response in TG mice linked to increased PC death. A. Representative flow cytometry plots of B220^+^ CD138^+^ plasmablasts (PBs) and B220^-^ CD138^+^ plasma cells (PCs) in the spleen of unimmunized naïve mice (left) and quantification of PB and PC numbers (right) (*n*= 8 WT and 5 TG). B. Quantification of serum immunoglobulin (Ig) titers from unimmunized WT and TG mice (*n* = 6 WT and 5 TG). C. Quantification of TNP-specific IgM serum titers from WT and TG mice 7 days post-immunization with TNP-Ficoll (*n* = 6 WT and 6 TG). D. Quantification of NP-specific IgM and IgG1 serum titers from WT and TG mice 14 days post-immunization (*n* = 6 WT and 5 TG). NP-specific IgG1 serum titers were re-assessed 35 days post-immunization followed by re-immunization with NP-CGG. NP-specific IgG serum titers were quantified 7 days post re-immunization to assess the secondary immune response (*n*= 3 WT and 3 TG). E. Representative flow cytometry plots of the splenic PC population (boxed area) in mice infected with the Armstrong strain of LCMV 8 days post-infection (dpi) (left) and quantification of total splenic PC numbers (right) (*n* = 4 WT and 4 TG). F. Quantification of total IgG serum titers from LCMV infected mice 8 dpi. G. Quantification of LCMV-specific IgG serum titers from LCMV infected mice 8 dpi. H-J. Naïve splenic mouse B cells from WT and TG mice were labeled with CFSE and then stimulated with CD40L, IL-4, and IL-5. Cells were collected and analyzed at days 0, 3, 4, and 5 of stimulation (*n* = 5 WT and 4 TG). Representative flow cytometry plots of WT and TG PC differentiation. Boxed areas indicate the percentage of PCs (H). Representative flow cytometry plots showing number of cell divisions by TG and WT B cells at day 5 of stimulation by CFSE dye dilution assay (I). Live cells (DAPI^-^) were gated into early divisions, late divisions, and PC subpopulations and the number of cells for each quantified by flow cytometry (J). Quantification of caspase 3/7 activation in *in vitro* derived PCs (K) (*n* = 4 WT and 4 TG). Data information: In (A), data is presented as mean ± SEM. In (B-D,F,G,J,K), data are presented as mean ± SD. *P* values were determined by two-tailed, unpaired Student’s *t*-test, \**P ≤* 0.05*, **P ≤* 0.01*, ***P ≤* 0.001*, ****P ≤* 0.0001.

We next performed ELISPOT assays to enumerate ASC frequencies. The color intensity and/or size of each spot within a well represents the amount of antibody secreted by each ASC. Consistent with a reduced number of splenic PCs in naïve mice with reduced Ig titers, splenic IgM-specific ASCs generated from TI immunizations were reduced over time in TG compared to WT mice (Fig EV2A and B). Of note, although there is no statistical difference in the number of ASCs at day 7, a difference in spot intensities, with blue spots being most intense, was detected between WT and TG ASCs (Fig EV2B). By day 14 however, only blue spots are left remaining, indicating a reduction in splenic ASCs in TG compared to WT mice (Fig EV2A and B). In ASCs generated from a TD response, there was also a dependence on time and spot intensity in ASC results. IgM-specific ASCs were reduced in number from the spleen of TG compared to WT mice (Fig EV2C and D). For IgG1-secreting ASCs however, although statistically similar in number between TG and WT mice, TG IgG1 ASCs exhibited less intense spots than WT IgG1 ASCs (Fig EV2C and D). For more mature ASCs, those representing PCs that migrated into the bone marrow, both IgM- and IgG1-secreting PCs were reduced in number and spot intensity in TG compared to WT mice (Fig EV2E and F). Interestingly, LLPCs are known to import more glucose and secrete more antibodies than SLPCs, linking antibody secretion levels to length of PC survival (Lam et al., 2016, Lam & Bhattacharya, 2018). The detection of spots with varying intensities indicated heterogeneous populations of ASCs with different levels of antibody secretion and potentially different levels of glucose import. Thus, the data show that CRTC2 inactivation and repression in PCs supports the function and maintenance of PCs, and by extension possibly the length of PC survival.

To determine whether sustained *Crtc2-AA* expression effects were ASC intrinsic, isolated naïve splenic B cells were first activated *in vitro* with CD40L, IL-4, and IL-5. A time course analyses of PC formation and/or persistence suggested that a reduction in the number of ASCs and impaired humoral immune responses could be cell intrinsic because TG PCs decreased over the course of stimulation, ending with a reduction from 37.3 to 18.4 percent PCs at day 5 (Fig 3H). The decrease in TG compared to WT PCs was not related to changes in B cell activation, proliferation, GC formation, or CSR during B cell differentiation (Fig EV3). Instead, we found that CRTC2 inactivation supported the survival of B cells at an advanced stage of differentiation, as the number of TG B cells were reduced in a division dependent manner compared to WT controls at day 5 of *in vitro* stimulation (Fig 3I). Quantification of cell numbers by flow cytometry gating on early divisions, late divisions, and PC subpopulations revealed a significant decrease in cell number only in the TG PC subpopulation compared to WT controls (Fig 3J). Flow cytometry analyses of *in vitro* derived and sorted PCs also revealed that TG PCs contained an increased level of activated, pro-apoptotic caspase 3/7 protein expression compared to WT PC controls (Fig 3K). This increase in cell death occurred at a differentiation stage when native *Crtc2* expression was robustly repressed in WT PCs but was sustained in an active form by the *Crtc2-AA* transgene in TG PCs (Fig EV1F). With the loss of *in vitro* generated PCs (Fig 3H and J) and a reduction in splenic and bone marrow ASCs over time (Fig EV2), TG PCs appeared to exhibit a cell intrinsic survival defect. Thus, physiologic CRTC2 inactivation and repression is required for PC survival in the spleen and bone marrow, directly affecting the generation of a robust, long-lived (Fig EV2) humoral immune response.

### CRTC2 regulates mature B cell mRNA transcription and splicing

To gain insight into reduced PC survival with sustained *Crtc2-AA* expression, we performed RNA-seq on naïve and sorted early division, late division, and PC TG and WT B cell subpopulations (Fig EV1E). Principle-component analysis (PCA) of the RNA-seq data showed that each replicate within a given subpopulation clustered together as expected, and that TG and WT B cell subsets were similar until modest separation occurred in the PC subpopulation (Fig 4A). In concordance, *Crtc2-AA* induced the largest number of transcript differences for TG versus WT cells at the PC stage, with 767 differentially expressed genes (p-adj < 0.05). Most differential genes were up-regulated (502/767, 65%), consistent with the established role of CRTCs as transcription coactivators (Altarejos & Montminy, 2011) (Fig 4B and Table EV1). By contrast, 38, 54, and 26 genes were differentially-expressed in naïve, early division, and late division subpopulations, respectively (p-adj < 0.05) (Fig 4B and Table EV1). Of the differentially-expressed genes within the PC subpopulation, 192 had >|2|-fold (Log_2_ fold > |1|) changes in expression (Fig 4C). Key regulators of B cell fate and survival, such *as Prdm1*, *Irf4*, and *Mcl1* showed no significant differences in TG compared to WT PCs (Peperzak, Vikstrom et al., 2013, Shapiro-Shelef, Lin et al., 2005, Shi et al., 2015, Tellier, Shi et al., 2016) (Fig 4D). However, *Cd28*, a costimulatory protein expressed in activated T cells and ASCs, was repressed 2.6-fold in TG compared to WT PCs (Delogu, Schebesta et al., 2006, Linsley & Ledbetter, 1993) (Fig 4D). CD28 functions as a pro-survival factor for LLPCs in the bone marrow but is dispensable for the survival of SLPCs in the spleen (Rozanski, Arens et al., 2011). A recent report found that differential expression of *Lcp2*, which encodes the CD28 adaptor protein SLP76, linked CD28 signaling in LLPCs to increased *Irf4* expression and an IRF4-mediated increase in mitochondrial mass and oxidative phosphorylation. This increase in metabolic fitness was proposed as a mechanism for LLPC survival in the bone marrow niche (Utley, Chavel et al., 2020). However, a previous study failed to identify *Lcp2* as differentially expressed between ASCs and follicular B cells (Shi et al., 2015) and our own RNA-seq analysis showed no significant difference in *Lcp2* expression between TG PCs and WT PCs. Also, gene set variation analysis (GSVA) of IRF4 and BLIMP1 target genes that are important for PC survival and function (Tellier et al., 2016) showed no differences between TG and WT subpopulations (Fig 4E). Furthermore, the unfolded protein response (UPR), and canonical early activation, GC, and ASC gene signatures were also found to be similar between TG and WT subpopulations (Fig 4E). Finally, analysis of all *Ig* transcripts between TG and WT B cell subpopulations were similar, indicating that the observed reduction in Ig secretion in TG ASCs by ELISPOT assay was not from changes in transcript levels (Table EV1).

**Figure 4.**
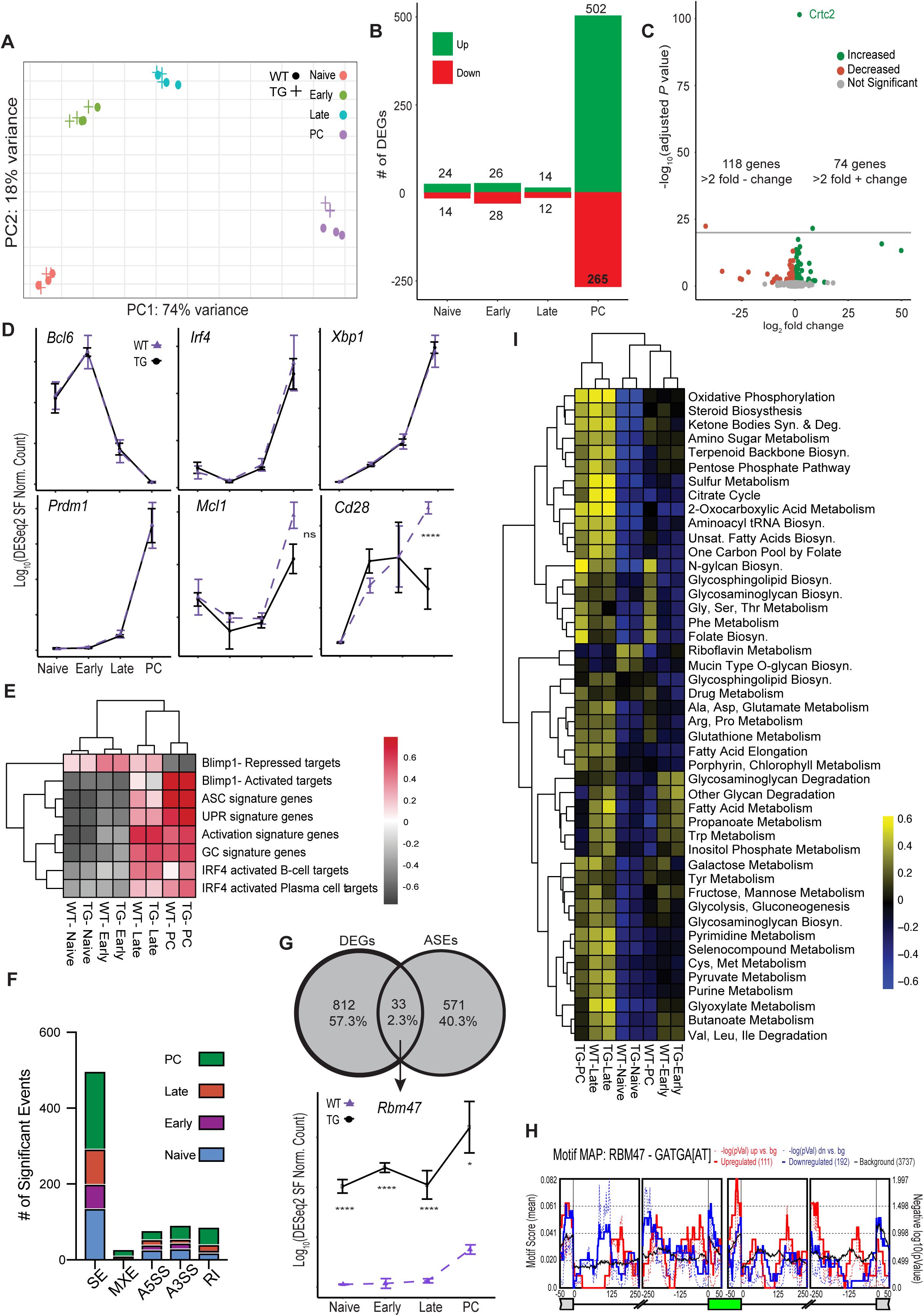
CRTC2 regulates mRNA expression and alternative splicing in mature B cells. A. Principal component analysis of bulk transcriptomes derived from CD138 and CFSE dye-dilution FACS sorted populations described in Fig EV1F. Points represent replicates of *n* = 3 independent experiments for WT and TG pairs. PC1 and PC2 scores are plotted on the x- and y-axes, respectively. B. List of differentially expressed genes (DEGs) between WT and TG cells at each stage of PC fate specification derived from bulk transcriptomics. Numbers represent the number of up-regulated and down-regulated DEGs in each subpopulation. C. Volcano plot of WT and TG cell fold change differences during the PC stage of differentiation. Green dots indicate genes up-regulated in the TG population during the PC stage of differentiation, whereas red dots indicate down-regulated genes. The adjusted *P* value cutoff represents values < 0.05 calculated using the Wald test following DESeq2 normalization. D. Stage-course expression plots for transcripts of canonical B cell differentiation, PC formation, and PC survival markers *Bcl6, Irf4, Xbp1, Prdm1, Mcl1, and Cd28*. Data represent log_2_ transformed DESeq2 normalized counts. E. Gene set variation analysis (GSVA) results for *Blimp1* target, *Irf4* target, unfolded protein response (UPR) signature, and canonical early activation/GC/ASC formation signatures in WT and TG cell populations at each assessed stage of differentiation. Data represent *n* = 3 independent experiments. F. mRNA alternative splicing events (detected by rMATS 4.0.2) at each stage of PC differentiation. Events are split into skipped exon (SE), mutually exclusive exon (MXE), alternative 5’ (A5SS) or 3’ (A3SS) splice sites, and retained intron (RI) events. Events are detected at an inclusion level difference of 0.10 and an FDR < 0.05 at all stages of differentiation between WT and TG cells. G. Venn diagram displaying the intersection of all DEGs from and all mRNA alternative splicing events (ASEs) (top) and stage-course expression plot of *Rbm47* (bottom). H. Enrichment map of RBM47 binding motif in all SE events in the PC subpopulation analyzed by rMAPS. The green box represents an average exon and the flanking exons are represented in grey boxes. Numbers below represent nucleotide positions. Solid red and blue lines represent the mean RBM47 motif score at each nucleotide position. Dotted lines represents -log10 (pValues) of motif scores between regulated and background exons and shows significant enrichment for the RBM47 binding motif. I. GSVA results for metabolic signature genes derived from the KEGG database. Results represent metabolic gene sets passing a Benjamini Hochberg adjusted P value threshold < 0.01. Data represent the mean of *n* = 3 independent experiments. Data information: Adjusted *P* values were determined by Wald test, \**P ≤* 0.05, \*\*\*\**P ≤* 0.0001

To identify candidate biological processes influenced by CRTC2 in PCs, we examined differentially expressed genes in the PC subpopulation by KEGG pathway analysis. The top three significantly enriched pathways were DNA replication, cell cycle, and spliceosome genes (Table EV1). We therefore assessed the cell cycle status of *in vitro* differentiated PCs at day 5 and found they were predominantly in the G2/M phase of the cell cycle with no significant differences between TG and WT PCs. Of note, recent reports have shown a role for alternative splicing in B cell fate (Chang, Li et al., 2015, Diaz-Munoz, Bell et al., 2015, Litzler, Zahn et al., 2019, Monzon-Casanova, Screen et al., 2018), which prompted an assessment of CRTC2 in alternative RNA splicing during B cell differentiation. Although CRTCs do not contain conserved RNA binding domains, one *in vitro* study in non-immune cells suggested that CRTCs may regulate mRNA splicing (Altarejos & Montminy, 2011, Amelio, Caputi et al., 2009). It is unknown, however, whether CRTC2 impacts mRNA splicing in activated B cells maturing into PCs. Analysis of the RNA-seq data revealed significant alterations (by >10%) in the splicing of 604 transcripts in TG compared to WT cells, most of which favored skipped exons (SE) and occurred in the PC subpopulation (Fig 4F and Table EV2). Of note, only 33 (2.3%) gene transcripts were altered by both expression and splicing between TG and WT B cells, indicating that aberrant splicing is not responsible for the majority of gene expression changes in TG cells (Fig. 4G and Table EV2). Interestingly, *Rbm47,* an RNA binding protein that modulates alternative splicing and mRNA abundance in the context of cancer (Sakurai, Isogaya et al., 2016, Vanharanta, Marney et al., 2014) was significantly altered in both expression and splicing, resulting in the enrichment of a processed isoform of *Rbm47* which has no known function (Fig 4G and Table EV2). Of note, knockdown of *Rbm47* in a lung cancer cell line resulted in an increase in oxidative phosphorylation (Sakurai et al., 2016). In addition, the RBM47 binding motif was the most enriched motif in SE events in the PC subpopulation (Fig 4H), suggesting that CRTC2 regulation of RBM47 could regulate PC physiology. However, similar to expression changes observed by RNA-seq, most of the changes in transcript splicing were small in magnitude and revealed no clear functional or biological enrichment for reductionist testing to help explain a decrease in TG PC survival (Table EV2).

Since the length of PC survival was previously linked to unique metabolic programs (Lam et al., 2016), we assessed whether CRTC2, with established roles in energy homeostasis and mitochondrial metabolism (Burkewitz, Morantte et al., 2015, Han, Kwon et al., 2020, Wu, Huang et al., 2006), impacted the expression of genes that participate in cellular metabolism. Surprisingly, a selective GSVA for metabolism regulating genes revealed that TG PCs show a metabolic gene expression profile similar to late dividing WT and TG PC precursors that clustered distinctly from WT PCs (Fig 4I). In addition, TG PCs showed an enrichment for the oxidative phosphorylation gene set, which is noteworthy because a switch from glycolysis to oxidative phosphorylation is required for ASC differentiation (Price, Patterson et al., 2018), while genes significantly dysregulated by CRTC2 constitutive activation, *Cd28* and *Rbm47*, have been shown to affect oxidative phosphorylation. Thus, CRTC2 independently regulated gene expression and mRNA splicing patterns during B cell differentiation and its repression mediated the fidelity of these processes in PCs, including genes that regulate PC metabolism.

### *Crtc2* repression increases oxidative metabolism

To determine whether the *Crtc2-AA* induced transcript changes between TG and WT PCs was sufficient to alter cellular metabolism, we measured the cellular oxygen consumption rate (OCR) and extracellular acidification rate (ECAR) as surrogates for mitochondrial respiration and glycolysis using a Seahorse Extracellular Flux Analyzer. OCR measurements on *in vitro* derived and sorted CD138^+^ PCs revealed similar basal respiration with reduced maximal respiration and reduced spare respiratory capacity (SRC) for TG PCs compared to WT controls (Fig 5A-D). SRC is utilized by cells under conditions of increased work or stress and has been correlated with cell longevity (D’Souza & Bhattacharya, 2019, van der Windt, Everts et al., 2012). In addition, suppressed pyruvate generation and/or mitochondrial pyruvate import is known to reduce SRC in LLPCs and leads to a progressive loss of LLPCs in the bone marrow (Lam et al., 2016). ECAR measurements in TG PCs were also lower compared to WT controls, suggesting less available pyruvate for glycolysis or oxidative metabolism (Fig 5E). Analysis of the mitochondrial membrane potential (MMP) by tetramethylrhodamine ethyl ester (TMRE) staining revealed a reduction in MMP in TG PCs (Fig 5F), which is consistent with reduced respiration (Fig 5A). The enrichment of the oxidative phosphorylation gene set in TG PCs (Fig 4I) may be part of a known compensatory mechanism in which cells with reduced mitochondrial function increase the expression of nuclear and mitochondrial genes in an attempt to recover oxidative metabolism (Reinecke, Smeitink et al., 2009). Also, changes in total mitochondrial mass can impact mitochondrial functioning, including cellular respiration (Liesa & Shirihai, 2013). We therefore utilized MitoTracker Green staining to assess mitochondrial mass and found no significant differences between TG and WT PCs (Fig 5G). The data suggest that reduced respiratory capacity in TG PCs is not due to differing amounts of mitochondria, but may be due to differences in the ability to import and/or utilize respiratory substrates.

**Figure 5.**
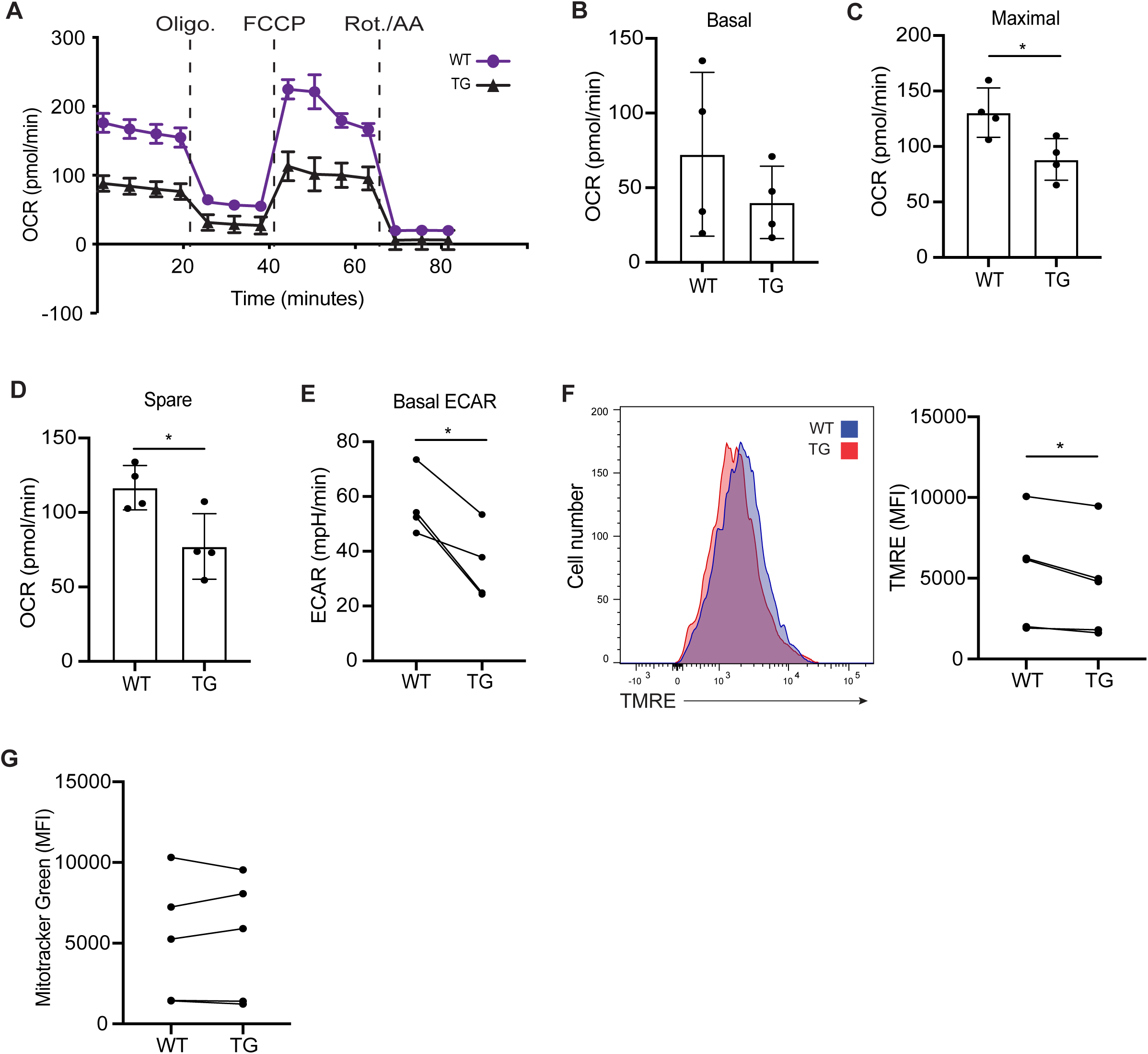
*Crtc2* repression increases oxidative metabolism. A-E. *In vitro* differentiated cells were sorted for CD138^+^ PCs at day 5 and metabolism was measured using the Seahorse Extracellular Flux Analyzer. Each point represents the mean ± SD of *n* = 4 independent experiments. Representative extracellular flux analysis of WT and TG PCs. Oxygen consumption rate (OCR) before and after treatment with indicated pharmacological inhibitors (A). Quantification of basal oxygen consumption rate (B). Quantification of maximal oxygen consumption rate (C). Quantification of spare respiratory capacity (D). Quantification of basal extracellular acidification rate (ECAR) from extracellular flux assay in A (E). F-G. *In vitro* differentiated cells were sorted for CD138^+^ PCs at day 5 and were stained with the indicated dyes. The mean fluorescence intensity (MFI) was assessed by flow cytometry (*n* = 4 WT and 4 TG). Representative flow cytometry plot of TMRE fluorescence (left) and quantification of the MFIs for TMRE staining (right) (F). Quantification of the MFIs for Mitotracker Green staining (G). Data information: *P* values were determined by two-tailed, unpaired Student’s *t*-test (C,D), and paired Student’s *t*-test (E), \**P ≤* 0.05.

To test this idea, we cultured *in vitro* derived and sorted PCs with ^13^C_6_-labeled glucose to track the flux of glucose-derived carbon atoms. Although increased PC lifespan has been correlated with increased glucose import (Lam et al., 2016, Lam et al., 2018), there was an equivalent amount of glucose imported into TG and WT PCs (Fig EV4A). Labeled carbons from glucose were predominantly incorporated into 3-phosphoglycerate (3PG) and lactate with only modest incorporation into TCA cycle metabolites (Fig EV4B). The fraction of glycolytic and TCA cycle metabolites generated from ^13^C_6_-labeled glucose showed no significant differences between TG and WT PCs (Fig EV4B). However, there was a trend in TG PCs in which ^13^C-labeled TCA cycle metabolites represented a smaller fraction of the total metabolites compared to WT PCs, suggesting that *Crtc2-AA* may modestly reduce glucose utilization. In addition, steady-state metabolite analyses of TG and WT PCs identified glycerol-3-phosphate (G3P), acetylcholine (Ac-choline), and guanine monophosphate (GMP) as decreased in TG compared to WT PCs (Fig EV4C). G3P is noteworthy as a metabolite that delivers cytosolic reducing equivalents into the mitochondria through a G3P shuttle to help fuel oxidative phosphorylation. However, the lack of large differences in glucose utilization, total metabolites levels, and metabolic processes (Table EV3) between TG and WT PCs suggests that *Crtc2* repression influences gene expression and alternative splicing programs that broadly and modestly increase cellular metabolism in PCs.

### *Crtc2* repression increases PC longevity

We identified reductions in survival (Fig 3H-K), antibody secretion (Fig 3 and Fig EV2), glycolysis (Fig 5E), maximal respiration (Fig 5A and C), and SRC (Fig 5A and D) in TG compared to WT ASCs. Therefore, we suspected that WT SLPCs could express more *Crtc2* transcripts than WT LLPCs because of strong similarities between SLPCs and the *Crtc2-AA* mouse model. Consistent with this prediction, independent RNA-seq profiling of PC subsets with different half-lives (Lam & Bhattacharya, 2018) showed highest *Crtc2* expression in the shortest half-life PC subset and the lowest expression in the longest half-life PC subset (Fig 6A). This data mining difference in *Crtc2* mRNA transcripts clearly trended but was not statistically significant between PC subsets. However, the PC subsets analyzed were not pure populations because of current technical limitations in identifying and isolating heterogeneous PC populations (Lam & Bhattacharya, 2018), and thus may account for the lack of statistical significance in *Crtc2* expression despite this clear trend.

**Figure 6.**
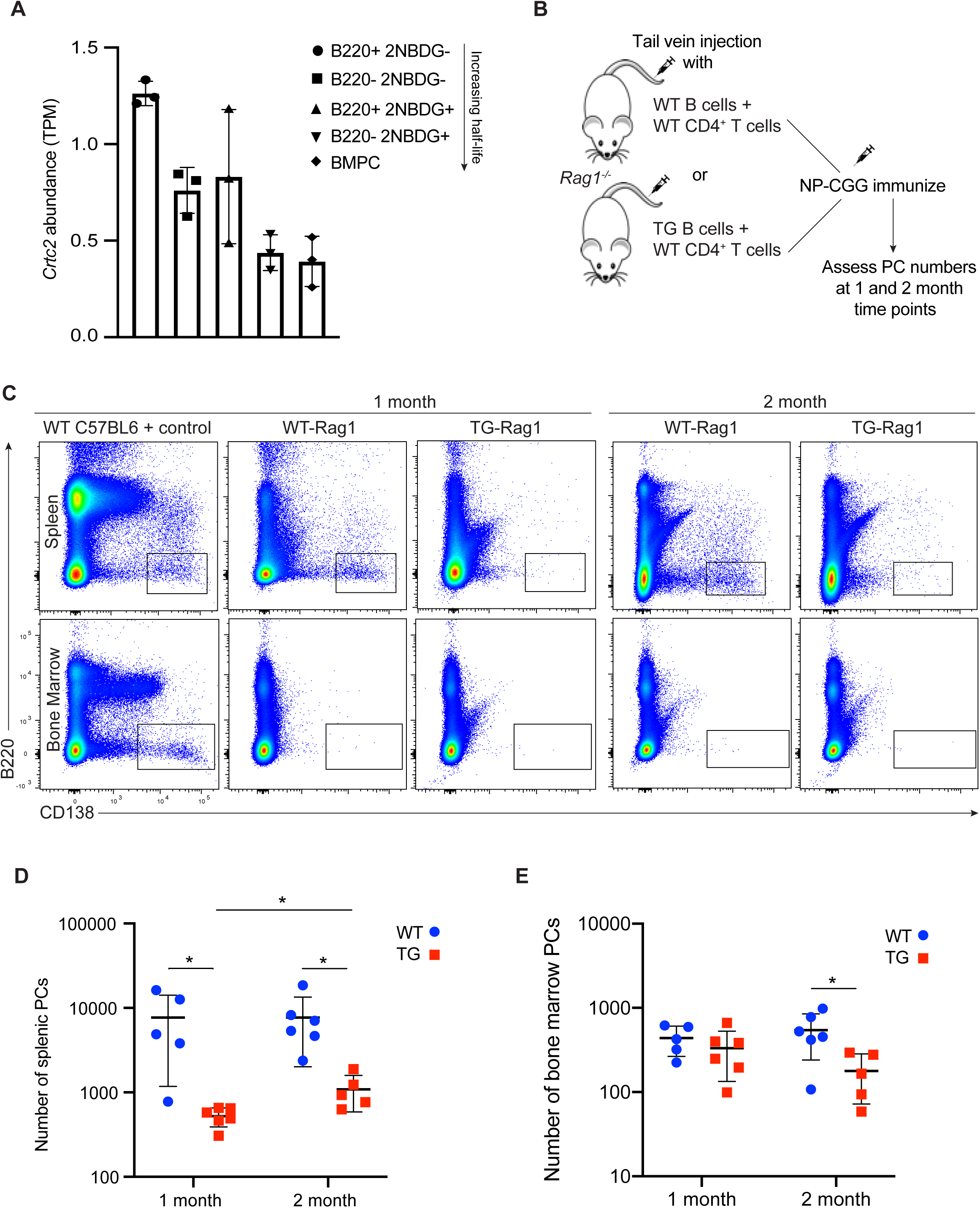
*Crtc2* repression supports increased PC longevity. A. Abundance of *Crtc2* transcripts, measured in transcripts per million (TPM) in sorted PC subsets from the spleen and bone marrow. RNA-seq data from Lam et al., 2018, was re-analyzed to determine transcript abundance. The half-life of sorted PC subsets increases from left to right as previously determined by Lam et al., 2018. Data represents the mean ± SD of *n* = 3 independent experiments. B. Experimental outline for adoptive transfer-immunization experiment. Two independent experiments were performed for each time point. Each symbol represents an individual mouse and the data are cumulative from two independent experiments for each time point (*n* = 5 WT and 6 TG at 1 month, 6 WT and 5 TG at 2 month). C. Representative flow cytometry plots of splenic and bone marrow PC populations (boxed areas) in WT-Rag1 and TG-Rag1 mice. WT C57BL6 mice was used as a positive control to set gating parameters. D. Quantification of total splenic PC numbers. E. Quantification of total bone marrow PC numbers. Data information: In (D,E), data are presented as mean ± SD. *P* values were determined by two-tailed, unpaired Student’s *t*-test, \**P ≤* 0.05.

To examine the effect of failed CRTC2 inactivation on PC longevity and to further examine cell intrinsic versus extrinsic effects, we performed an adoptive transfer-immunization experiment (Weisel, Zuccarino-Catania et al., 2016). We transferred equal numbers of naïve CD43^-^ TG or WT B cells with WT CD4^+^ T cells into lymphocyte-deficient *Rag1^-/-^* recipient mice. Transfer recipients with WT B cells (WT-Rag1) and TG B cells (TG-Rag1) were immunized one day post-transfer with NP-CGG to initiate a TD immune response for a comparative assessment of the number of PCs over time (Fig 6B). In the spleen, the number of TG PCs at 1- and 2-month time points were reduced compared to WT controls (Fig 6C and D). Over the 2 month time-course, there was an unanticipated 2-fold increase in TG PCs whereas the number of WT PCs remained unchanged (Fig 6D). This result could be from experimental variability between TG-Rag1 cohorts, variability in immunogen administration, and/or a modest difference in CD4^+^ T cell help (Fig EV5). Analysis of the bone marrow revealed few PCs overall, with a significant decrease in the number of TG PCs compared to WT PCs at the 2 month time point, and a 46% decrease in TG PCs compared to a 24% increase in WT PCs between 1 and 2 month assay points (Fig 6C and E). Overall, these results confirm that reduced PC survival in TG mice is B cell intrinsic and caused by sustained expression and activity of CRTC2. Both RNA-seq data mining analyses and *in vivo* adoptive transfer studies support a role for CRTC2 inactivation in PC longevity.

## Discussion

Our results demonstrate the requirement for physiologic *Crtc2* repression in PCs in order to sustain a humoral immune response. Elevated expression of a *Crtc2-AA* transgene at this stage in differentiation altered metabolism and reduced the survival of PCs, resulting in humoral immune deficits against model antigens and an acute viral infection. Differentiation into PBs, however, was unaffected by CRTC2-AA, as TG mice had a similar number of splenic PBs as WT mice. In addition, similar numbers and frequencies of ASCs generated by immunizations and *in vitro* differentiation, respectively, at early time points uncovered a reduction in survival from failed CRTC2 inactivation within the more mature PC population. Our results apply to both TI and TD immune responses, which can both generate LLPCs that typically migrate to and localize within the bone marrow (Bortnick, Chernova et al., 2012).

The metabolic requirements of PCs differ from those of activated B cells and differentiating PBs. Instead of relying on energy and biosynthetic pathways for nucleotide biosynthesis and organelle biogenesis during rapid replication and cell expansion, PCs switch their metabolism to mainly support antibody production^7^. Aligned with this shift in metabolic requirements*, Crtc2* repression and CRTC2 inactivation in PCs is necessary for the fidelity of PC gene expression and mRNA alternative-splicing programs. When these programs were altered by sustained CRTC2-AA activity, TG PCs adopted a SLPC-like phenotype (Lam & Bhattacharya, 2018), showing reductions in antibody secretion, glycolysis, oxidative phosphorylation, and SRC. The reduction in PC longevity did not depend on differences in glucose import, but rather appeared more systemic with small, broad reductions in metabolite utilization in TG compared to WT PCs. Carbon tracing studies identified a modest trend toward lower ^13^C-label incorporation into glycolytic and TCA cycle metabolites and a reduction in total G3P abundance in TG compared to WT PCs. This lack of large, specific changes in glucose utilization and total metabolite abundance suggests that CRTC2 inactivation may help fine tune cellular metabolism based on levels of *Crtc2* repression, perhaps similar to the broad regulatory role played by the MYC transcription factor family, as an example. In lymphocytes, MYC binds thousands of genes and acts to broadly amplify the expression of bound genes rather than functioning as an on/off switch (Nie, Hu et al., 2012). By an imperfect analogy, we identified CRTC2 bound to more than 4,500 genes in naïve WT mouse B cells by ChIP-seq (unpublished observation), many of which were not predicted as CREB target genes, in agreement with prior results that showed CRTC proteins interact with additional bZIP family transcription factors (Escoubas, Silva-Garcia et al., 2017). In TG PCs, large changes in expression of specific genes were few, but small changes in expression did occur for numerous genes between TG and WT PCs. This more modest, rheostat-like change in the expression of multiple genes could underlie a modest scaled difference in metabolism between TG and WT PCs. Importantly, prior carbon tracing studies of glutamine in human LLPCs identified a robust contribution to glutamate and aspartate, but no labeling of TCA metabolites citrate or aconitate (Lam et al., 2018). Here, we did not identify differences in total glutamate or aspartate abundance between TG and WT PCs, suggesting that glutamine tracing may yield mainly uninformative results and further support for the idea that the extent of CRTC2 inactivation in PCs could help to set the overall level of metabolism, or metabolic fitness, of these cells.

As an alternative and perhaps not mutually exclusive idea, TG PCs showed differential splicing of RNA transcripts associated with electron transport chain (ETC) activity and ETC complex formation compared to WT PCs. These ETC protein-encoding transcripts included *Cep89*, *Ndufaf7*, and *Sirt3*. CEP89 is a known regulator of ETC Complex IV activity (van Bon, Oortveld et al., 2013), which transfers electrons to molecular oxygen, whereas NDUFAF7 and SIRT3 regulate ETC Complex I assembly and activity, respectively (Ahn, Kim et al., 2008, Zurita Rendon, Silva Neiva et al., 2014). Changes in the splicing of these and perhaps additional transcripts could underlie the reduced respiratory capacity of TG compared to WT PCs that even a compensatory elevation in the expression of oxidative phosphorylation genes could not overcome (Fig 4I). Furthermore, the splicing of these and other transcripts with SE events may be regulated in part by CRTC2 control of expression and alternative splicing of *Rbm47*. In this scenario, the extent of CRTC2 inactivation may have a more targeted effect on substrate utilization and ETC activity rather than, or in addition to, helping to set overall cellular metabolism in PCs.

The metabolism of LLPCs differs from SLPCs (Lam et al., 2016, Lam et al., 2018, Utley et al., 2020), although the mechanism(s) underlying differences in lifespan remain largely unknown. Due to multiple similarities between SLPCs and PCs generated by the *Crtc2-AA* transgenic model, we hypothesized that PC longevity might correlate with levels of *Crtc2* repression, which analysis of an independent RNA-seq dataset supported with a clear trend between lower *Crtc2* transcript levels and increased PC lifespan (Fig 6A). Based on our current and prior studies that demonstrated CRTC2 inactivation through B cell antigen receptor (BCR) engagement followed by ATM, LKB1, and AMPK family member signaling (Kuraishy, French et al., 2007, Sherman et al., 2010), we provide a testable model for additional studies in which the level of CRTC2 activity in PCs is determined by the BCR-antigen interaction and relayed strength of signaling. The level of CRTC2 activity, mediated by the extent of transcript repression and protein inactivation, would help establish levels of antibody secretion, glycolysis, oxidative phosphorylation, and SRC that support metabolic fitness and survival through effects on gene expression and alternative transcript splicing (Fig 7). Thus, PCs with higher affinity antibodies would show a survival advantage over PCs with lower affinity antibodies. In addition, this model is consistent with data from additional biological systems that linked CRTC family member proteins with lifespan regulation. For example, AMPK extended the lifespan of *C. elegans* through CRTC1-dependent remodeling of mitochondrial metabolism (Burkewitz et al., 2015). Additionally, IgA secreting PCs, which show a high turnover rate (Pabst, 2012), may be too short-lived to be impacted by CRTC2 regulation of PC survival, which could help to explain why IgA titers were unaffected in unimmunized *Crtc2-AA* TG mice (Fig 3B).

**Figure 7.**
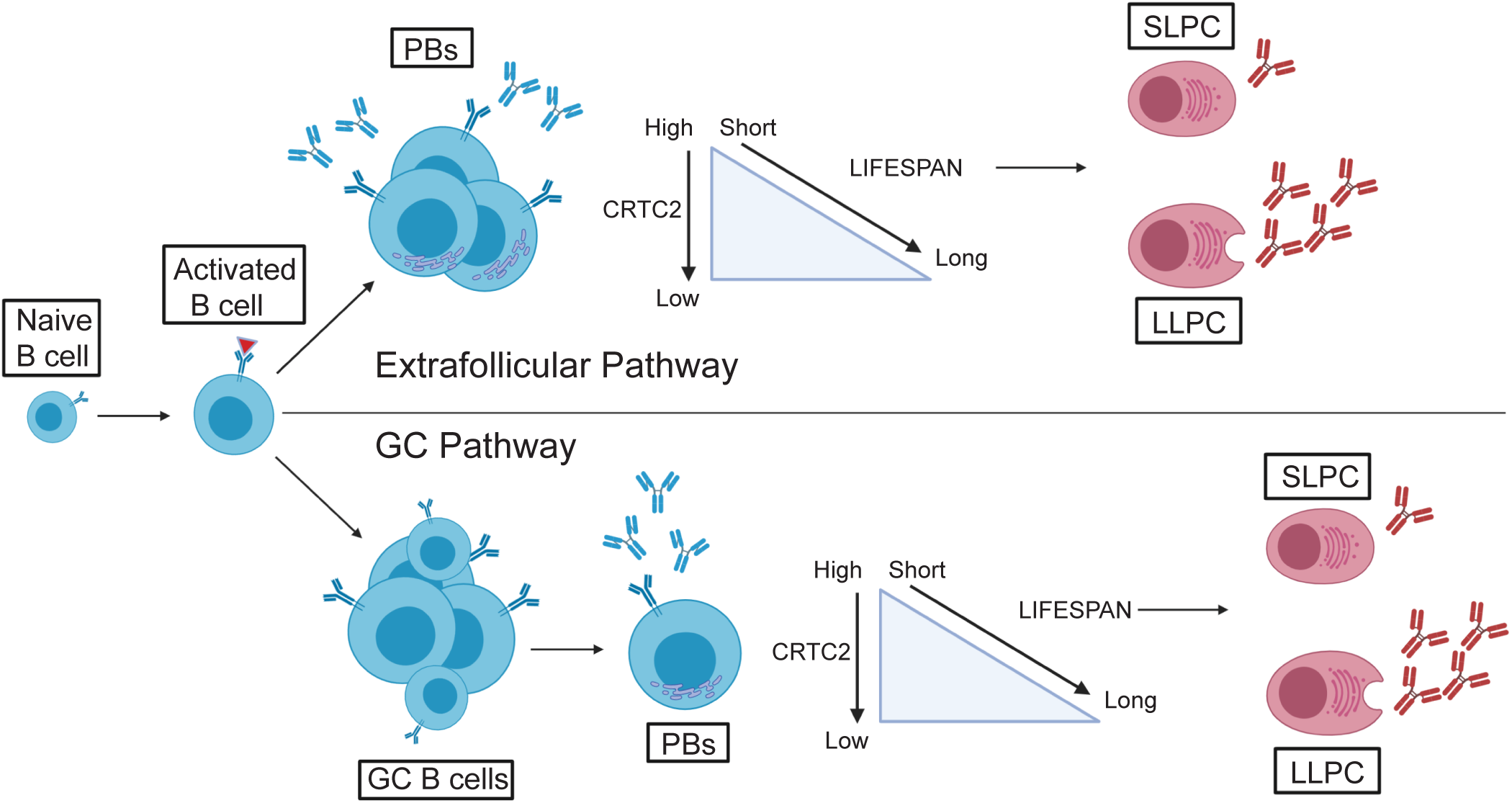
Proposed model for CRTC2 regulation of PC survival and longevity. Details are provided in the text.

Previously, we showed that sustained CRTC2-AA expression in isolated and stimulated human B cells blocked ASC differentiation by maintaining activated B cells in a GC B cell-like state. The difference in outcome between those results and our report here may relate to differential biological functions of CRTC2 at different stages of B cell development rather than inconsistencies between studies. This is similar to BCL6 and IRF4, which have different functions depending on stage of B cell differentiation (Duy, Yu et al., 2010, Ochiai, Maienschein-Cline et al., 2013, Phan & Dalla-Favera, 2004). We speculate that when CRTC2-AA expression is enforced in early activated human B cells, ASC differentiation is blocked, whereas enforced CRTC2-AA expression in mouse PCs results in altered metabolism and shortened survival. In support of this idea, disruption of upstream signaling components in the CRTC2 inactivation pathway show effects consistent with early CRTC2 dysregulation. Patients harboring mutations *in ATM* exhibit defective antibody responses (Staples, McDermott et al., 2008) and mouse B cells lacking LKB1 fail to generate ASCs *in vitro* (Walsh, Waters et al., 2015). Identifying direct CRTC2-bound genes may help elucidate potential stage-specific targets, although this may prove challenging as CRTCs bind and co-activate multiple bZIP transcription factors in addition to CREB (Escoubas et al., 2017). In addition, CRTC family member proteins may have redundant functions, making stage-specific CRTC2 deletion studies difficult to interpret. For example, a single allele of *Crtc2* or *Crtc3* is capable of rescuing the viability of *Crtc2/*3 double knockout mice (Kim, Hedrick et al., 2017), potentially explaining why in nearly a decade, no reports have emerged on defects in the B cell compartment of *Crtc2* knockout mice (Hernandez, Chang et al., 2015, Le Lay, Tuteja et al., 2009, Wang, Inoue et al., 2010). Despite these confounding biological issues, identifying CRTC2 targets can uncover an additional layer of CRTC2 gene regulation. For example, although CRTC2 is a known coactivator, our prior study identified numerous genes that were induced by CRTC2 inactivation (Sherman et al., 2010). This suggests that CRTC2, directly or indirectly, could repress the expression of CRTC2 bound genes by preventing access to additional transcription or chromatin remodeling factors. The Montminy lab Online CREB database predicts that *Cd28* regulatory regions contain CREB response element binding sites and therefore may be bound by CRTC2. Although speculative, our unpublished data identified a CRTC2 interaction with specific chromatin modifiers, making this mode of regulation a possibility worth further exploration.

Since its discovery as a key regulator of the CREB transcription factor family nearly two decades ago, the CRTC protein family has been linked to numerous human pathologies including multiple cancer types, aging, and metabolic disorders (Escoubas et al., 2017). Here, we show that inactivation of CRTC2 by transcript repression and/or protein phosphorylation is required to ensure long-term PC survival and function. This study also points to potential therapeutic benefits of targeting and manipulating the level of CRTC2 activity in PCs, to either amplify a beneficial, or inhibit a harmful, humoral immune response.

## Materials and Methods

### Mice and animal procedures

*Crtc2-AA* transgenic mice were derived by pronuclear injection of C57BL/6 mouse embryos at the UCLA and UCI transgenic mouse facilities. Briefly, the mouse *Crtc2* coding sequence with introduced S171A and S275A alterations was cloned into a *B29* minimal promoter, *IgH* intronic enhancer (*Eμ*) expression plasmid (Hoyer et al., 2002). The plasmid was injected into the pronucleus of mouse embryos and then implanted in a surrogate dam for gestation. F0 generation pups were selected for transgene integration through tail biopsy and PCR analysis and three founder lines were identified. For *in vivo* experiments, mice were immunized by intraperitoneal administration of 2 x 10^8^ sheep red blood cells (Colorado Serum Company), 25 ug of 2,4,6-Trinitrophenyl (TNP)-aminoethylcarboxymethyl-ficoll (AECM-Ficoll) (Biosearch Technologies), or 50 ug of alum-precipitated 4-Hydroxy-3-nitrophenylacetyl (NP)-chicken gamma globulin (CGG) (Biosearch Technologies). For adoptive transfer experiments, 10 week old female *Rag1^-/-^* mice (Jackson Laboratory, stock 002216) were injected via tail vein with 1×10^7 sorted CD43^-^ WT or TG naïve B cells and 5×10^6 sorted WT CD4^+^ T cells. Mice were immunized one day post adoptive transfer with 50 ug of NP-CGG. CD4^+^ T cells were isolated from spleen cells by immunomagnetic negative selection (Miltenyi Biotec). For viral infections, mice were inoculated intravenously with 2 x 10^6^ plaque forming units of the LCMV Armstrong strain. Virus stocks were prepared and viral titers were quantified as described (Elsaesser, Sauer et al., 2009). Mouse experiments were conducted on mixed-sex mice between 8-12 weeks of age. No animals were excluded due to a lack of responsiveness to immunization or infection. Randomization, but not experimental blinding to sample identity was used for these studies. All animals were housed in a pathogen-free animal facility at UCLA and all procedures were performed with approval from the UCLA Animal Research Committee (#1998-113-63).

### Enzyme-linked immunosorbent (ELISA) & enzyme-linked immunospot (ELISPOT) assays

Serum Ig and TNP- and NP-specific antibodies were detected by ELISA as described (Waters, Ahsan et al., 2019). TNP- and NP-specific ASCs were detected by ELISPOT assay. Briefly, MultiScreen HTS-HA filter plates (Millipore) for ELISPOT assays were prepared with 35% ethanol and then coated overnight at 4 °C with TNP-BSA or NP-BSA (Biosearch Technologies). Plates were washed and subsequently blocked at 37 °C with complete culture media prior to plating and culturing splenocytes or bone marrow cells for 18 h at 37 °C. Plates were washed with PBS-Tween (0.05%) and double-distilled water to remove cells before being incubated with alkaline phosphatase-conjugated anti-mouse IgM or anti-mouse IgG1 (Southern Biotech) for 2 h at room temperature. Plates were then washed with PBS-Tween (0.05%) and PBS before developing with BCIP/NBT blue substrate (Southern Biotech). Spots were imaged with an ImmunoSpot Analyzer (Cellular Technologies Ltd.) and quantified using ImageJ.

### B cell isolation and cell culture

Whole spleens were mechanically disassociated through a 40 μm filter followed by red blood cell lysis with ammonium-chloride-potassium lysing buffer. Naïve mature B cells were isolated from spleen cells by immunomagnetic negative selection with anti-CD43 (Miltenyi Biotec). B cells were cultured in RPMI-1640 medium (Corning) supplemented with 10% FBS (Gibco), 1% penicillin/streptomycin (Corning), 1% MEM non-essential amino acids (Gibco), 1% sodium-pyruvate (Corning), 1% L-glutamine (Gibco), and β-mercaptoethanol (50 μM). B cells were stimulated with 1 ug/ml CD40L (BD Pharmingen) and 25 ng/ml IL-4 (R&D Systems) or with 0.1 ug/ml CD40L, 10 ng/ml IL-4, and 5 ng/ml IL-5 (R&D Systems) to monitor B cell processes during B cell differentiation or to monitor differentiation with increased plasma cell (PC) frequency, respectively (Shi et al., 2015). For isolation of *in vitro* generated PCs used for Seahorse Extracellular Flux Analyzer assays, cultures stimulated for 5 days with CD40L, IL-4, and IL-5 were enriched for live cells using a dead cell removal kit (Miltenyi Biotec) followed by CD138-APC (BD Pharmingen) staining. CD138^+^ PCs were enriched by positive immunomagnetic enrichment of APC (Miltenyi Biotec). For Caspase 3/7, Mitrotracker Green, and TMRE staining experiments, *in vitro* generated PCs were isolated by immunomagnetic positive selection with anti-CD138 (Miltenyi Biotec).

### Flow cytometry

Single cell suspensions were stained with conjugated antibodies and data were obtained on a BD LSR II (BD Biosciences) and analyzed with FlowJo software (Treestar). Cell sorting was performed on a BD FACSAria (BD Biosciences). B and T cell populations, B cell class-switching, and B cell activation were analyzed with anti-B220, anti-IgM, anti-IgG1, anti-CD4, anti-CD8, anti-CD19, anti-CD21, anti-CD23, anti-CD25, anti-CD43, anti-CD69, anti-CD86, anti-CD93, anti-CD95(FAS) anti-CD138, and anti-GL7. Cell populations analyzed were as follows: pro-B cells, B220^+^IgM^-^CD43^+^; pre-B cells, B220^int^IgM^-^CD43^-^; immature B cells, B220^int^IgM^+^CD43^-^; mature B cells, B220^hi^IgM^+^CD43^-^; follicular B cells, B220^+^CD19^+^CD93^-^CD23^+^CD21^-/int^; and marginal zone B cells, B220^+^CD19^+^CD93^-^CD23^-^CD21^+^; germinal center B cells, B220^+^GL7^+^CD95^+^; plasmablast (PB), B220^+^CD138^+^; plasma cell (PC), B220^-^CD138^+^; double negative T cells (DN), CD19^-^CD4^-^CD8^-^; double positive T cells (DP), CD19^-^CD4^+^CD8^+^; CD4 T cells, CD19^-^CD4^+^; and CD8 T cells, CD19^-^CD8^+^. Representative flow cytometry gating strategies and reagents are provided in **Appendix Fig S1** and **Appendix Table S1** respectively. DAPI was used for assessment of cell viability and *in vitro* B cell proliferation was assessed by flow cytometry after labeling of isolated CD43^-^ naive mature B cells with CellTrace carboxyfluorescein succinimidyl ester (CFSE) (Invitrogen). *In vitro derived* PCs were stained with CellEvent Caspase 3/7-Green (ThermoFisher), tetrametylrhodamine, ethyl ester (TMRE) (Abcam), and Mitotracker Greeen (ThermoFisher) to assess caspase 3/7 activity, mitochondrial membrane potential, and mitochondrial mass by flow cytometry respectively. The number of donor derived cells in the spleen and bone marrow of adoptively transferred mice were calculated based on flow cytometric analysis of stained cells described above.

### Protein separation and immunoblot analysis

Total whole cell extracts were prepared by incubating cells in lysis buffer containing 50 mM Tris HCl pH 7.4, 100 mM NaCl, 1 mM EDTA, and 1% Triton X-100 supplemented with protease and phosphatase inhibitors (Sigma Aldrich). Cell extracts were quantified and loaded onto polyacrylamide SDS gels before separating by gel electrophoresis and transferring onto nitrocellulose membranes. ATM, pATM (S1981), pATM/ATR substrate, CRTC2, pCRTC2 (S171), HDAC1, α-actin, and β-tubulin proteins were detected by immunoblot analysis with antibodies listed in **Appendix Table S1**. Protein bands were visualized by chemiluminescence or IRDye detection at 680 or 800 nm using an autoradiograph film developer or the Odyssey Fc imaging system (LI-COR).

### RNA extraction and RT-PCR assays

Total RNA for quantitative RT-PCR assays were extracted using Trizol reagent (Invitrogen). Assays were performed with Kapa SYBR Fast qPCR master mix (Roche) and mRNA expression levels of *Crtc2* and *Prdm1* were quantified with standard curves and results were normalized to the expression of *Rps18*. RT-PCR primer sequences are provided in **Appendix Table S2**. RNA for sequencing were extracted using the RNeasy Mini Kit (Qiagen) and RNase-free DNase (Qiagen). All samples showed an A260/280 ratio > 1.99. Prior to library preparation, quality control of the RNA was performed using the Advanced Analytical Technologies Fragment Analyzer (Advanced Analytical Technologies, now Agilent Technologies) and analyzed using PROSize 2.0.0.51 software. RNA Quality Numbers were computed per sample between 8.7 and 10, indicating intact total RNA per sample prior to library preparation.

### RNA-seq library preparation

Strand-specific ribosomal RNA depleted RNA-seq libraries were prepared from 1 µg of total RNA using the KAPA Stranded RNA-seq Kit with Ribo-Erase (Roche). Samples were prepared in at least triplicate for analysis. Briefly, rRNA was depleted from total RNA samples, the remaining RNA was heat fragmented, and strand-specific cDNA was synthesized using a first strand random priming and second strand dUTP incorporation approach. Fragments were then A-tailed, adapters were ligated, and libraries were amplified using high-fidelity PCR. All libraries were prepared in technical duplicates per sample and resulting raw sequencing reads merged for downstream alignment and analysis. Libraries were paired-end sequenced at 2×150 bp on an Illumina NovaSeq 6000.

### RNA-seq data processing

Raw sequencing runs were filtered for low quality reads and adapter contamination using FastQC (http://www.bioinformatics.babraham.ac.uk/projects/fastqc), SeqTK (https://github.com/lh3/seqtk), and Cutadapt (Martin, 2011). Filtered reads were quantified and quasi-mapped to the *Mus musculus* Gencode M17 (GRCm38.p6) reference transcriptome using the alignment-free transcript level quantifier Salmon v.0.9.1 (Harrow, Frankish et al., 2012, Mudge & Harrow, 2015, Patro, Duggal et al., 2017). The resulting estimated transcript counts were summarized into normalized gene level transcripts per million (TPM) and estimated count matrices using R (v. 3.4.0) Bioconductor (v. 3.5) package tximport (v. 1.4) (Soneson, Love et al., 2015).

The resulting sample gene count matrix was normalized and analyzed for differential gene expression using R (v. 3.4.0) Bioconductor (v. 3.5) package DESeq2 v1.16.0 (Huber, Carey et al., 2015, Love, Huber et al., 2014). Significance testing was performed using the Wald test, testing for the significance of deviance in a full design “Batch + GenotypeStage”, modeling the genotype effect at each division and unstimulated timepoint while accounting for batch variance between matched stimulation replicates. Resulting *P* values were adjusted for multiple testing using the Benjamini-Hochberg procedure (Benjamini & Hochberg, 1995). Differentially expressed genes (DEGs) were filtered using an adjusted false discovery rate (FDR) *P* value < 0.05 and an absolute log_2_(Fold Change) > 0.5 in either the stimulated or unstimulated paired conditions.

Volcano plots were made with ggplot2 as described above, using the DESeq2 output adjusted *P* values and log_2_-fold changes per TG/WT comparison at each timepoint. Division course kinetic plots were made using ggplot2 with DESeq2 size factor normalized counts using function plotCounts(). Gene expression heat maps were prepared using pheatmap() with row Z-scores calculated as the variance stabilized transform (VST) subtracted by the row mean. PCA was plotted using VST values with function plotPCA() with parameter “ntop = 100000”.

Pathway-level signature gene set enrichment analysis was performed using R Bioconductor package GSVA v1.26.0 function gsva() with parameters “method = gsva, rnaseq = FALSE, abs.ranking = FALSE, min.sz = 5, max.sz = 500” using a log_2_(TPM + 1) transformed gene expression matrix(Hanzelmann, Castelo et al., 2013). GSVA pathway enrichment scores per sample were extracted and assessed for significance using R Bioconductor package limma v3.34.9, as described above except with a Benjamini-Hochberg adjusted *P* value threshold = 0.01. Differentiation signature gene sets was obtained for 24 hr CD40L plus IL-4 activated B cells (Shi et al., 2015, Stein, Dutting et al., 2017), GC B cells (Gloury, Zotos et al., 2016, Shi et al., 2015), ASCs (Gloury et al., 2016, Shi et al., 2015), BLIMP1 activated and repressed targets (Minnich, Tagoh et al., 2016), and IRF4 activated-B cell and plasma cell targets (Ochiai et al., 2013).

Alternative splicing analysis was performed using rMATS v4.0.2 (Shen, Park et al., 2014). Alternative splicing events were selected using an inclusion level difference greater than 0.1 with an FDR < 0.05 for any event comparison between Naive, Early, Late, and PC stages. RNA binding motif analysis was performed using rMAPS (Park, Jung et al., 2016) using JCEC counts and standard web server settings. Gene ontology (GO) term overrepresentation analysis was performed using the combined ASE lists at any event comparison, with R packages and parameters as previous described (Shimada, Ahsan et al., 2018).

Lists of transcript/gene-level expression values and differentiation signature GSVA results are included in supplemental excel file Table EV1. Lists of alternative splicing events and associated GO terms are provided in supplemental excel file Table EV2.

### Transgene insertion analysis

Identification of the transgene insertion site following founder selection was performed using BBMap (v.June 11, 2018) (BBtools - sourceforge.net/projects/bbmap/). Following WGS of the TG and WT litter-matched mates, sequences were first aligned to the transgene sequence cassette using parameters “bbmap.sh in1=jh_wt[tg]_fixedmate_1.fastq.gz in2=jh_wt[tg]_fixedmate_2.fastq.gz outm=eitherMapped_jh_wt[tg]_fixedmate.fq ref=insertionsequence.fasta”. The resulting aligned sequences were then remapped to the Mus musculus mm10 reference genome (NCBI38/mm10, December 2011) to identify sites spanning transgene and native genome sequence, using parameters “bbmap.sh in=eitherMapped_jh_wt[tg]_fixedmate.fq ref=mm10.fasta out=genomeMapped_jh_wt[tg]_fixedmate.sam bs=bs.sh”. The resulting bam files were converted into bigWig format and visualized using IGV v2.3.81(Robinson, Thorvaldsdottir et al., 2011).

### Metabolomics

CD138^+^ PCs were sorted on day 4 of *in vitro* differentiation and grown for 18 hrs in glucose free 1640 RPMI (Thermo) with dialyzed FBS supplemented with 2 g/L [U-^13^C_6_] glucose (Cambridge Isotope Laboratories). Metabolites were extracted and measured using Ultra High Performance Liquid Chromatography Mass Spectrometry (UHPLC-MS). Briefly, cells were pelleted by centrifugation and rinsed with cold 150 mM ammonium acetate (pH 7.3). Following centrifugation, metabolites were extracted from cell pellets by adding 1 ml of cold 80% MeOH in water, 10 nmol of D/L-norvaline, followed by rigorous vortexing and centrifugation. The supernatants were transferred into glass vials and cell pellets were extracted once more with 200 ul of 80% MeOH. The supernatants from the second extraction were collected into the same glass vial and metabolites were dried under vacuum and resuspended in 70% acetonitrile. Mass spectrometry analysis was performed as previously described (Waters et al., 2019). Data analysis, including principal components analysis (PCA) and clustering, was performed using the statistical language R v4.0.0 and Bioconductor v3.11.0 packages. Metabolite abundance was normalized per µg of protein content per metabolite extraction, and metabolites not detected were set to zero. Metabolite normalized amounts were scaled and centered into Z-scores for relative comparison using R base function scale() with parameters “scale = TRUE, center = TRUE”. Volcano plots were prepared using R package ggplot2 v. 3.3.0; adjusted P values were calculated using R base functions t.test() and p.adjust() with parameter “method = ‘BH’” using corrected isotopomer distribution (MID) values.

PCA was performed using R packages FactoMineR v2.3 and factoextra v1.0.7. Normalized metabolite amounts were standardized using a log_2_(normalized amounts + 1) transformation, and PC scores computed with function PCA() using parameters “scale.unit = TRUE, ncp = 10, graph = FALSE”. PCA individual score plots were displayed using function fviz_pca(). PCA variable loadings plots were generated using function fviz_pca_var(), extracting metabolite scores for contributions to the top ten principal components, strength of representation on the factor map (cosine2) and variable coordinates indicating Pearson correlation coefficient *r* of each metabolite to the top ten principal components. For whole relative amounts PCA, variable loadings were classified into k=4 separate clusters using k-means clustering in function kmeans() using parameters “set.seed(123), centers = 4, nstart 25”. K-means clustering of glutamine and glucose MID tracing variable loadings were perform using k=2 clusters with parameters “set.seed(123), centers = 2, nstart = 25”.

Pathway-level relative amounts metabolite set variation analysis (MSVA) was performed using R Bioconductor package GSVA v1.36.3 (Hanzelmann et al., 2013). Metabolite normalized relative abundances were standardized using a log_2_(normalized amounts + 1) transformation, and metabolites per sample were converted to a pathways per sample matrix using function gsva() with parameters “method = gsva, rnaseq = FALSE, abs.ranking = FALSE, min.sz = 5, max.sz = 500”. GSVA pathway enrichment scores were then extracted and significance testing between conditions was calculated using R Bioconductor package limma v3.44.3, fitting a linear model to each metabolite and assessing differences in normalized abundance using an empirical Bayes moderated F-statistic with an adjusted *P* value threshold of 0.05, using the Benjamini-Hochberg false discovery rate of 0.05 (Benjamini & Hochberg, 1995, Ritchie, Phipson et al., 2015). Pathway metabolite sets were constructed using the KEGG Compound Database and derived from the existing Metabolite Pathway Enrichment Analysis (MPEA) toolbox (Kanehisa, Goto et al., 2012, Kankainen, Gopalacharyulu et al., 2011). Assessment of differentially produced metabolites was calculated between genotype (TG/WT, all samples), using R base function t.test, and corrected for multiple testing using p.adjust() with parameter “method = ‘BH’”. Metabolites were threshold at *P* < 0.05 or adjusted *P* < 0.05. Metabolite relative amounts, isotopomer distribution values, % labeling fractional contribution, and MSVA scores are included in Table EV3.

### Statistical and reproducibility

All metabolomics and transcriptomics statistical analyses are described in the above methods. Values represent mean ± SD or SEM. Data were analyzed with Prism 8 (GraphPad) (Figs 1-6 and Figs EV1-5) or R (Fig 4 and Fig EV1). Parametric data were analyzed using two-tailed, unpaired and paired Student’s *t*-tests, or 2-way ANOVA with Bonferroni correction for multiple comparisons. Transcriptomic volcano and kinetic time-course expression plots were analyzed using DESeq2 Wald tests with Benjamini-Hochberg FDR correction for multiple comparisons. For all data sets, P ≤ 0.05 was considered significant. *P ≤ 0.05, **P ≤ 0.01, ***P ≤ 0.001, ****P ≤0.0001.

## Data availability

All raw RNA-Seq and WGS reads, transcript abundance values, and processed gene count matrices are in submission to the NCBI Gene Expression Omnibus (GEO). Accession numbers will be provided upon request prior to publication. All RNA-seq, alternative-splicing, and metabolomics datasets have been provided as supplemental material in this study (Tables EV1-3). All custom code used for analyses are available on Atlassian BitBucket at https://bitbucket.org/ahsanfasih/crtc2bcell/src. All other relevant data are available from the corresponding author on request.

## Acknowledgements

We thank the UCLA Flow Cytometry Core, Immuno/Biospot Core, Mitochondria and Metabolism Core, and Metabolomics Center for technical support. We thank Drs. Douglas Black, Jessica Fowler, Nicole Walsh, Rani Najdi, Tara TeSlaa, Laura Jimenez, and Alexander Patananan, along with Alexander Sercel and Vivian Lu for technical assistance and helpful discussions. Supported by NIH awards R01CA90571, R01CA156674, R01GM073981, R21CA227480, and P30CA016042, and the Air Force Office of Scientific Research FA9550-15-1-0406.

## Author contributions

JSH, FMA, EM-R, PDP, and MAT designed experiments; JSH, FMA, EM-R, PDP, M-SL, TLN, DGB, JG, and KD performed experiments and/or analyzed data; FMA carried out computational analysis; KRN provided resources for RNA and whole genome sequencing; JSH and MAT wrote the manuscript with input and editing from FMA, PDP, DGB, and KRN.

## Expanded View Figure and Table Legends

**Figure EV1 - Transgenic founder lines, transgene expression, and transgene integration analysis.**

A. Schematic of the *Crtc2-AA* transgenic construct used to generate TG mouse founder lines.

B. *Crtc2-AA* transgene expression in bone marrow cells of each TG founder line by quantitative RT-PCR analysis (*n*= 3 for each founder line).

C. Quantification of absolute cell numbers from bone marrow B cell subpopulations in TG founder lines F2 and F3 compared to WT controls (*n* = 6 WT and 3 each F2, F3 TG).

D. Paired-end whole genome sequencing (WGS) of the F1 TG founder mouse to identify transgene integration site(s). Genome browser tracks depict mapping of reads to the mouse genome with integration sites determined by accumulation of reads that partially align to both the genome and the transgene insertion cassette. Data are plotted relative to WT WGS results analyzed in the same pipeline.

E. Representative flow cytometry plot of naïve splenic B cells stimulated with CD40L, IL-4, and IL-5 used for cell sorting and RNA-seq. At day 5 of stimulation, cells were sorted into early division, late division, and PC subpopulations as indicated by boxed areas based on CFSE dye dilution assay and CD138 staining.

F. *Crtc2* mRNA expression from sorted cell populations as outlined in E. Expression represented as normalized counts for 3 independent RNA-seq experiments of WT and TG pairs. Data information: In (B,C,F), data are presented as mean ± SD. *P* values were determined by two-tailed, unpaired Student’s *t*-test (**B,C**), adjusted *P* values were determined by Wald test (**F**), \**P ≤ 0.05*, *\*\*P ≤* 0.01, \*\*\**P ≤* 0.001, \*\*\*\**P ≤* 0.0001.

**Figure EV2 - ELISPOT images documenting reduction in ASC numbers with PC maturation.**

A. Quantification of TNP-specific IgM ASCs from the spleen 7 and 14 days post-immunization with TNP-Ficoll by ELISPOT (*n* = 3 WT and 3 TG for each 7 and 14 day experiment).

B. Representative images of ELISPOT wells in triplicate revealing IgM secreting ASCs at 7 days (left) and at 14 days (right) post TNP-Ficoll immunizations in WT and TG splenic cells.

C. Quantification of NP-specific IgM and IgG1 ASCs from the spleen 14 days post-immunization with NP-CGG plus alum by ELISPOT (*n* = 4 WT and 4 TG).

D. Representative images of ELISPOT wells in triplicate revealing IgM and IgG1 secreting ASCs at 14 days post NP-CGG immunizations in alum for WT and TG splenic cells.

E. Quantification of NP-specific IgM and IgG1 ASCs from the bone marrow 14 days post-immunization with NP-CGG plus alum by ELISPOT (*n* = 4 WT and 4 TG).

F. Representative images of ELISPOT wells in triplicate revealing IgM and IgG1 secreting ASCs 14 days post NP-CGG immunizations in alum for WT and TG bone marrow cells.

Data information: In (A,C,E), data are presented as mean ± SD. *P* values were determined by two-tailed, unpaired Student’s *t*-test, \**P ≤* 0.05*, **P ≤* 0.01.

**Figure EV3 - Unaffected B cell activation, proliferation, GC formation, and CSR in TG B cells.**

(A-C and E) Naïve splenic mouse B cells from WT and TG mice were stimulated with CD40L and IL-4 (*n* = 5 WT and 4 TG). Representative flow cytometry plots depicting activation marker expression at day 3 of stimulation (top) and quantification of activation marker expression by mean fluorescence intensity (MFI) (bottom) (A). Representative flow cytometry plots depicting cell division measured using CFSE dye dilution assay at day 3 of stimulation (B).

Representative flow cytometry plots depicting differentiation of GC-like B cells at day 3 of stimulation (left) and quantification of the percentage of GC-like B cells (right) (C).

D. Representative flow cytometry plots depicting differentiation of GC B cells (left) and quantification of the percentage and absolute number of GC B cells in the spleen of mice 12 days post SRBC immunization (*n* = 6 WT and 5 TG).

E. Representative flow cytometry plots depicting class switch recombination (CSR) to the IgG1 isotype at day 3 of stimulation (left) and quantification of the percentage of CSR (right).

Data information: In (A,C,D,E), data are presented as mean ± SD.

**Figure EV4 - Stable isotope ^13^C_6_-glucose tracing in TG and WT PCs.**

A. Mass isotopomer distribution of glucose.

B. Fractional contribution of ^13^C_6_-labeled metabolites from [U-^13^C_6_] glucose after 18 h quantified by UHPLC-MS.

C. Volcano plot distribution of differentially abundant UHPLC-MS identified metabolites. *P* values represents FDR uncorrected values.

Data information: Data from glucose tracing represent *n* = 3 independent experiments.

**Figure EV5 - Adoptive transfer-immunization experimental controls.**

A. Representative flow cytometry plots of the population of transferred B cells in the spleen (boxed areas).

B. Representative flow cytometry plots of the population of transferred CD4^+^ T cells in the spleen (boxed areas).

C. Quantification of the total number of B cells.

D. Quantification of the total number of CD4^+^ T cells.

Data information: In (C,D) data are presented as mean ± SD of *n* = 5 WT and 6 TG at 1 month and *n* = 6 WT and 5 TG at 2 month.

**Table EV1 -** List of transcript and gene-level expression values, GO term overrepresentation results, and signature differentiation pathway scores from RNA-seq datasets for WT and *Crtc2-AA* TG B cells during the differentiation stage-course.

**Table EV2** List of mRNA alternative splicing events, differential expression overlaps, and GO term overrepresentation analysis of significantly spliced genes.

**Table EV3**. Metabolomics analysis of CRTC2 AA TG and WT CD138^+^ PCs. List of relative amounts, mass isotopomer distributions, fractional contribution labeling, and GSVA results.

