## Appendix for "CRTC2 regulates plasma cell metabolism and survival"

**Appendix Figure S1  p. 2**

**Appendix Table S1   p. 4**

**Appendix Table S2   p. 5**

### Appendix Figure S1

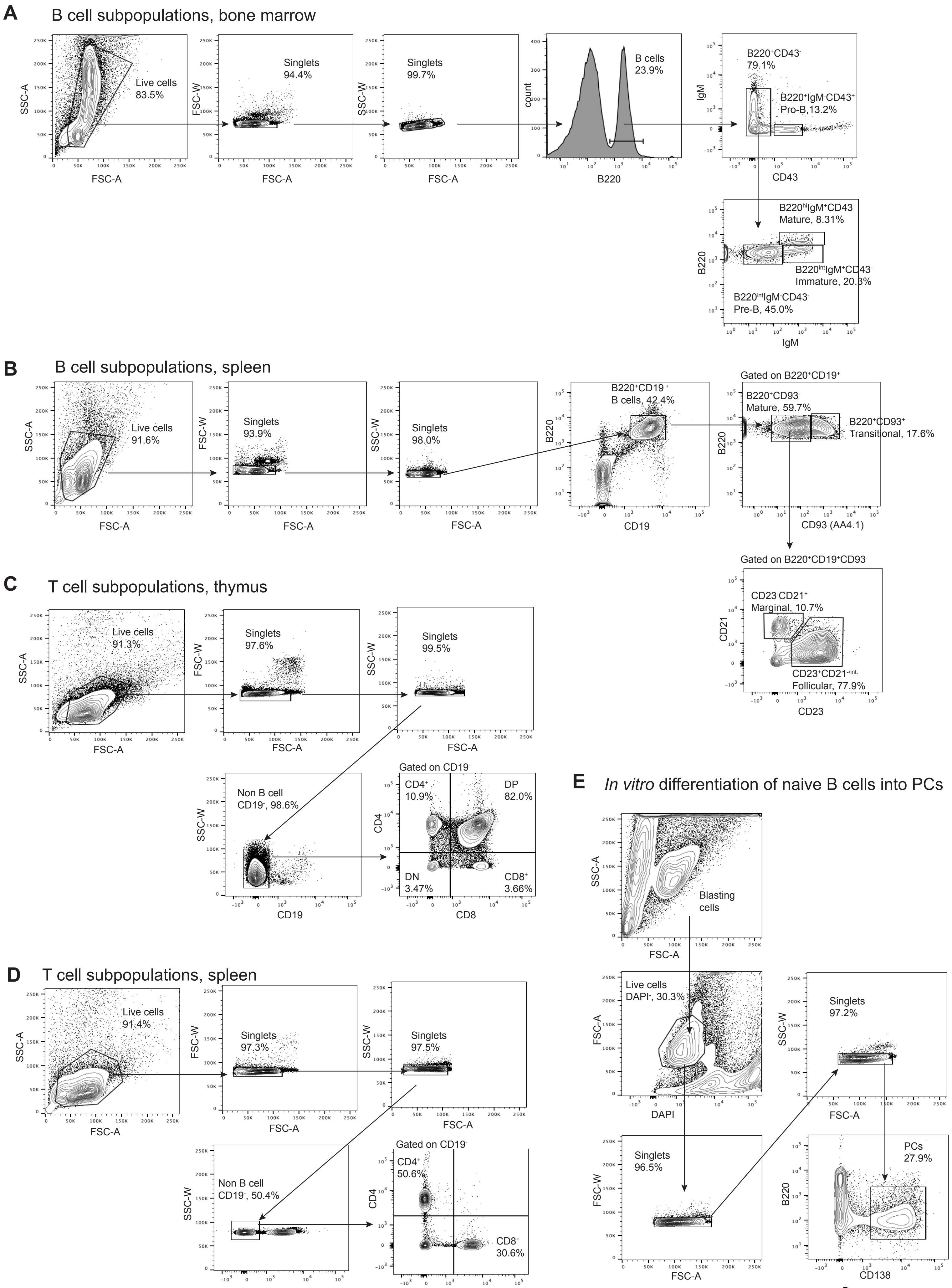

**Appendix Figure S1 - Representative gating strategy for flow cytometry analysis.**

A. Representative gating strategy for analyzing the B cell subpopulations in the bone marrow.

B. Representative gating strategy for analyzing the B cell subpopulations in the spleen.

C. Representative gating strategy for analyzing the T cell subpopulations in the thymus.

D. Representative gating strategy for analyzing the T cell subpopulations in the spleen.

E. Representative gating strategy for analyzing the *in vitro* differentiation of naïve splenic B cells into PCs with either CD40L and IL-4 or CD40L, IL-4, and IL-5 stimulations.

**Appendix Table S1 - List of antibodies used in this study.**

| Protein Detected | Source | Conjugate | Application | Product # |
|---|---|---|---|---|
| Actin | rabbit | unlabeled | WB | A2066 |
| ATM | mouse | unlabeled | WB | sc-23921 |
| ATM (phospho-S1981) | rabbit | unlabeled | WB | AF1655 |
| ATM/ATR substrate ab. (phospho-Ser/Thr) | rabbit | unlabeled | WB | 2851 |
| β-tubulin | mouse | unlabeled | WB | T4026 |
| CRTC2 | rabbit | unlabeled | WB | PA5-34547 |
| CRTC2 (phospho-S171) | rabbit | unlabeled | WB | ab203187 |
| HDAC1 | mouse | unlabeled | WB | sc-81598 |
| LKB1 | mouse | unlabeled | WB | sc-32245 |
| rabbit IgG | donkey | HRP | WB | 711-035-152 |
| rabbiit IgG | donkey | IRDye 680 | WB | 926-68073 |
| mouse IgG | goat | IRDye 800 | WB | 925-32210 |
| B220 | rat | APC | Flow Cytom. | 561880 |
| B220 | rat | Alexa Fluor 700 | Flow Cytom. | 56-0452-82 |
| B220 | rat | FITC | Flow Cytom. | 553087 |
| B220 | rat | PerCP-Cy5.5 | Flow Cytom. | 552771 |
| B220 | rat | PE-Cy7 | Flow Cytom. | 25-0452-81 |
| B220 | rat | PE-eFluor 610 | Flow Cytom. | 61-0452-82 |
| B220 | rat | PE | Flow Cytom. | 553090 |
| CD4 | rat | PE | Flow Cytom. | 553653 |
| CD8 | rat | eFluor 450 | Flow Cytom. | 48-0081-82 |
| CD16/CD32 | rat | unlabeled | Flow Cytom. | 553140 |
| CD19 | rat | APC Cy7 | Flow Cytom. | 115530 |
| CD21 | rat | FITC | Flow Cytom. | 11-0211-82 |
| CD23 | rat | PE | Flow Cytom. | 12-0232-81 |
| CD25 | rat | APC | Flow Cytom. | 17-0251-81 |
| CD43 | rat | PE | Flow Cytom. | 553271 |
| CD69 | rat | FITC | Flow Cytom. | 11-0691-81 |
| CD86 | rat | V450 | Flow Cytom. | 560450 |
| CD93 | rat | PE | Flow Cytom. | 554258 |
| CD138 | rat | PE/APC | Flow Cytom. | 553714/561705 |
| IgM | rat | PECy7 | Flow Cytom. | 25-5790-82 |
| IgG1 | rat | FITC | Flow Cytom. | 553443 |
| GL7 | rat | FITC | Flow Cytom. | 553666 |

| | | | | |
|---|---|---|---|---|
| CD40 | hamster | unlabeled | Cell activation | 553721 |
| mouse IgM | goat | AP | ELISPOT | 1020-04 |
| mouse IgG1 | goat | AP | ELISPOT | 1070-04 |
| mouse Ig | goat | unlabeled | ELISA | 5300-05 |
| mouse IgM | goat | HRP | ELISA | 5300-05 |
| mouse IgG1 | goat | HRP | ELISA | 5300-05 |
| mouse IgG2a | goat | HRP | ELISA | 5300-05 |
| mouse IgG2b | goat | HRP | ELISA | 5300-05 |
| mouse IgG3 | goat | HRP | ELISA | 5300-05 |
| mouse IgA | goat | HRP | ELISA | 5300-05 |
| mouse IgM | mouse | unlabeled | ELISA | 5300-01 |
| mouse IgG1 | mouse | unlabeled | ELISA | 5300-01 |
| mouse IgG2a | mouse | unlabeled | ELISA | 5300-01 |
| mouse IgG2b | mouse | unlabeled | ELISA | 5300-01 |
| mouse IgG3 | mouse | unlabeled | ELISA | 5300-01 |
| mouse IgA | mouse | unlabeled | ELISA | 5300-01 |

**Appendix Table S2 – List of oligonucleotide primers used in this study.**

**Oligonucleotides**

***Crtc2*** **(total)**

Fwd: AGA GTC TGG CTG GCG AAG

Rev: GCA GAC GGC AGT CTA AAC AA

***Crtc2*** **(transgene specific)**

Fwd: ACT CAT TCC GTA GTG ATC GGC TAC AG

Rev: AAG TAA GGT TCC TTC ACA AAG ATC CGG

***Prdm1***

Fwd: TGC GGA GAG GCT CCA CTA

Rev: TGG GTT GCT TTC CGT TTG

***Rps18***

Fwd: TTT GCG AGT ACT CAA CAC CAA

Rev: TTC CTC AAC ACC ACA TGA GC
